# Stimulus-Dependent Dopamine Dynamics from Locus Coeruleus Axons

**DOI:** 10.1101/2025.09.15.676390

**Authors:** Avi Matarasso, Itzel Rodriguez Reyes, Elena Seaholm, Ashritha Cheeyandira, Michael J Seibert, Suchitha Jagalur, Sean C. Piantadosi, Li Li, Yulong Li, David Weinshenker, Michael R. Bruchas

**Affiliations:** Department of Bioengineering, University of Washington, Seattle, WA, USA; Department of Anesthesiology and Pain Medicine, University of Washington, Seattle, WA, USA; Center for the Neurobiology of Addiction, Pain and Emotion; Departments of Anesthesiology, Pharmacology, and Bioengineering, University of Washington, Seattle, WA, USA; Department of Biology, University of Washington, Seattle, WA, USA; Department of Neuroscience, University of Washington, Seattle, WA, USA; Norcliffe Foundation Center for Integrative Brain Research, Seattle, WA, USA; State Key Laboratory of Membrane Biology, Peking University School of Life Sciences, Beijing, China; Department of Human Genetics, Emory University School of Medicine, Atlanta,GA, USA; Department of Pharmacology, Rice University, Houston, TX, USA

**Keywords:** locus coeruleus, dopamine, hippocampus, norepinephrine, optogenetics, fiber photometry, neuromodulation

## Abstract

Arousal is essential for survival, and maladaptive arousal processing leads to an inability to focus, anxiety-like behavior, and dysregulated affective states. Norepinephrine (NE) is known to regulate anxiety, arousal, and learning through locus coeruleus (LC) projections throughout the brain. Evidence for co-release of the NE precursor and neurotransmitter dopamine (DA) from LC neurons has been accumulating for years, yet definitive measures of DA release across regions, stimulus paradigms, and behaviors associated with the LC-NE system remain controversial. Here, we identified the physiological and behavioral properties that evoke DA release from LC axon terminals. Using concomitant approaches, we inhibited the LC and ventral tegmental area (VTA) to selectively isolate the contributions of LC-derived DA release. Together these findings establish the constraints by which LC neurons release DA in a modality-dependent manner.

## 1 INTRODUCTION

The locus coeruleus-norepinephrine (LC-NE) system consists of a widespread projecting network of neurons that modulate a range of behaviors like arousal (Berridge 2008, Carter et al. 2010), learning (Bouret and Sara 2004, Kaufman et al. 2020, Seo et al. 2021), and stress-induced negative affect (Giustino et al. 2020, McCall 2015, McCall et al. 2017). As an obligate precursor for NE, dopamine (DA) is also produced in LC neurons. However, because tyrosine hydroxylase (TH; converts tyrosine to L-DOPA), not dopamine *β*-hydroxylase (DBH; converts DA to NE), is rate-limiting for catecholamine synthesis under normal circumstances (**?**), LC neurons were classically thought to release NE but not DA. Yet, the potential for DA release from noradrenergic neurons has been discussed since the early 2000s. Initially, microdialysis was used to detect neuromodulator release with autoreceptor antagonists (which disinhibit NE or DA neuron activity), and revealed the potential for noradrenergic release of DA. For example, α2 adrenergic receptor (α2ARs) antagonists, but not a D2 dopamine receptor antagonist, upregulated DA release in the occipital cortex (OCC) and medial prefrontal cortex (mPFC), brain regions with very sparse dopaminergic innervation (Borgkvist et al. 2011, Devoto et al. 2003 2004 2005a). Furthermore, LC stimulation increased the amount of extracellular DA in mPFC and in OCC, which was diminished after local tetrodotoxin administration in mPFC or OCC (Devoto et al. 2005b). Overall, these data suggest that α2ARs on LC axonal terminals regulate release of DA (Devoto et al. 2005a 2003).

While DA could be evoked from noradrenergic neurons through α2AR antagonist administration, whether noradrenergic-sourced DA release affects behavior remained unclear. To investigate the role of LC-DA in behavior, studies employed LC terminal stimulation in the dorsal hippocampus (CA1) and found that spatial awareness and memory were enhanced (Kempadoo et al. 2016, Takeuchi et al. 2016). Infusion of high concentrations of D1 receptor antagonists in CA1 reversed these enhanced spatial memory effects, while blocking β-adrenergic receptors (βARs) had no impact.

These studies implicate LC-evoked DA release in behavior for the first time, but the durations and intensities employed in optogenetics experiments (Gálvez-Márquez et al. 2022, Kempadoo et al. 2016, Takeuchi et al. 2016) do not reflect the LC’s established neuronal activity during many of these tasks (Devilbiss and Waterhouse 2011, Kelberman et al. 2024). Pharmacological manipulations in these studies also do not preclude potential circuit level or non-selective effects of these manipulations. Moreover, a ventral tegmental area (VTA) to CA1 dopaminergic circuit controls persistent contextual memory (Lisman and Grace 2005) and contextual fear acquisition in the absence of noradrenergic signaling (Tsetsenis et al. 2021), affirming the need to determine the physiological and behavioral stimuli which evoke DA release from LC axons. There remains limited direct evidence for release of DA from LC terminals *in vivo* in response to appetitive, aversive, or neutral stimuli.

Previous studies detecting LC-evoked DA release utilized microdialysis, which while highly sensitive, lack temporal resolution (Kempadoo et al. 2016, Takeuchi et al. 2016). In contrast, recently developed genetically-encoded biosensors have risen to prominence for neuro-modulator detection due to their high temporal resolution and selectivity for target ligands. However, G protein-coupled receptors (GPCRs) have can have non-selective affinity for off-target neuromodulator-GPCRs of similar subclasses and derivatives, and off-target binding has been reported, specifically with NE and DA (López et al. 2024). So while GPCR-based sensors have become widely-used for understanding behavior at relevant time-scales, studies must carefully consider the local innervation of their targeted circuit for off-target neuromodulator release that may inadvertently activate endogenous receptors and sensors (López et al. 2024).

In this study, we consider the controversy surrounding LC-derived DA and how we may measure NE and DA dynamics in regions where there are some overlapping DA and NE inputs. Emerging evidence suggests LC neurons innervate regions with functional specificity and distinct subsets of neurons activate depending on behavioral state (Chandler et al. 2019, Dupret et al. 2010, Nakahara et al. 2004, Poe et al. 2020, Sinha et al. 1999, Uematsu et al. 2017). For example, during early extinction of conditioned fear, LC projections to basolateral amygdala (BLA) are preferentially active. By contrast, during late extinction when mice exhibit diminished fear behavior, LC projections to mPFC are active (Uematsu et al. 2017). The BLA is critical for cued fear (Campeau and Davis 1995, Davis and Whalen 2001, Fanselow and LeDoux 1999, Maren 2001, **?**, Phillips and LeDoux 1992) while CA1 is necessary in contextual fear conditioning (Anagnostaras et al. 1999, Maren et al. 1994, **?**, Phillips and LeDoux 1992), and both are also activated by salient stimuli (Barth et al. 2023, Liu et al. 2021, Schoenbaum et al. 1999, Shukla and Chattarji 2022, Uwano et al. 1995) and reward-associated cues (Ambroggi et al. 2008, Gauthier and Tank 2018, Martig and Mizumori 2011, Paton et al. 2006, Tye and Janak 2007, Uwano et al. 1995, Yun et al. 2023). Therefore, the BLA serves as a critical region alongside the hippocampus to further determine the specific nature of LC-evoked DA dynamics.

Since distinct subsets of LC neurons are activated in response to salient, aversive, and appetitive stimuli (Borodovitsyna et al. 2018, Chandler et al. 2019, Chen and Sara 2007, Curtis et al. 2012, de Medeiros et al. 2005, Grimm et al. 2024, McCall et al. 2017, Poe et al. 2020, Privitera et al. 2024, Silveira et al. 1993) in different behavior states, we hypothesized that DA release from the LC would be greater in BLA than in CA1 during both aversive and appetitive stimuli, as salient stimuli are often associated with BLA activity (Liu et al. 2021, Uwano et al. 1995) and the CA1 receives little dopaminergic input (Tsetsenis et al. 2021). We first established the selectivity constraints and demonstrated high selectivity of the sensors we used. We next demonstrated that we can evoke DA release from LC-CA1 terminals in 2-photon *ex vivo* slice imaging and then found that LC-evoked DA signal follows an increasing linear trend with frequency *in vivo* in CA1 and BLA. We next established the NE and DA responses to appetitive and aversive stimuli in the BLA and CA1. Finally, we used chemogenetics to demonstrate that following appetitive and aversive stimuli, the LC contributes more to endogenous DA release in the BLA than in CA1. Together, we report that LC releases DA with both projection and stimulus specificity. As diminished DBH has been associated with increased vulnerability to psychosis (Cubells et al. 2000, Meltzer et al. 1976, Yamamoto et al. 2003) and neurodegeneration in Parkinson’s disease (Lieberman et al. 1972, Nagatsu et al. 1982), characterizing the ratio of DA/NE release by the LC may help determine the neural mechanisms behind these diseases.

## 2 RESULTS

### GRAB_NE_ does not detect DA *in vivo* in response to biologically relevant photostimulation

Because activation of catecholamine sensors by the off-target catecholamine (DA in the case of GRAB_NE2m_) has been reported (López et al. 2024), it is important to test the limitations of these sensors *in vivo*. If ligand selectivity of each biosensor is not considered, then the results of the stimuli-dependent release could be misconstrued, across any region where dual NE and DA innervation is present. We compiled the most updated EC50s generated in vitro (i.e. the presumed KD due to lack of G-protein coupling) for DA and NE at each GPCR-based sensor for examining the difference in DA vs NE selectivity (Figure 1A) (Feng et al. 2024 2019, Patriarchi et al. 2018, Sun et al. 2020). Among the available sensors, the most selective DA sensor is GRAB_DA3m_, with an EC50 of 110nM for DA and 8μM for NE, representing an 80-fold difference in selectivity. The next most selective DA sensor at the time of this study was dLight1.3b, which has a 70-fold difference in selectivity, but also has a larger EC50 for NE than other sensors, at 59μM. GRAB_rDA2m_ has a 60-fold difference in selectivity, and similar EC50s for NE and DA compared to GRAB_DA3m_. Compared to dLight, the GRAB_DA3m_ sensors have a higher affinity for both the target catecholamine and for the non-targeted catecholamine. NE has less affinity at GRAB_NE2m_ than DA has at GRAB_DA3m_ sensors. Comparing on/off kinetics of these cate-cholamine biosensors is a powerful method of determining separability of recorded signals. The dLight1.3b sensor has a *τ* _on_ (10ms) 5-7 times faster than GRAB_DA3m_ (69ms) and GRAB_rDA2m_ (50ms) (Figure 1B). While they have a similar *τ* _on_ rate (50-69ms), the red GRAB_rDA2m_ sensor has *τ* _off_ (2.24s) 4 times slower than the GRAB_DA3m_ sensor (0.56s). Here, we use these kinetics to compare neuromodulator recordings during behavior and qualitatively explain the likely separability of simultaneously recorded signals.

**FIGURE 1.**
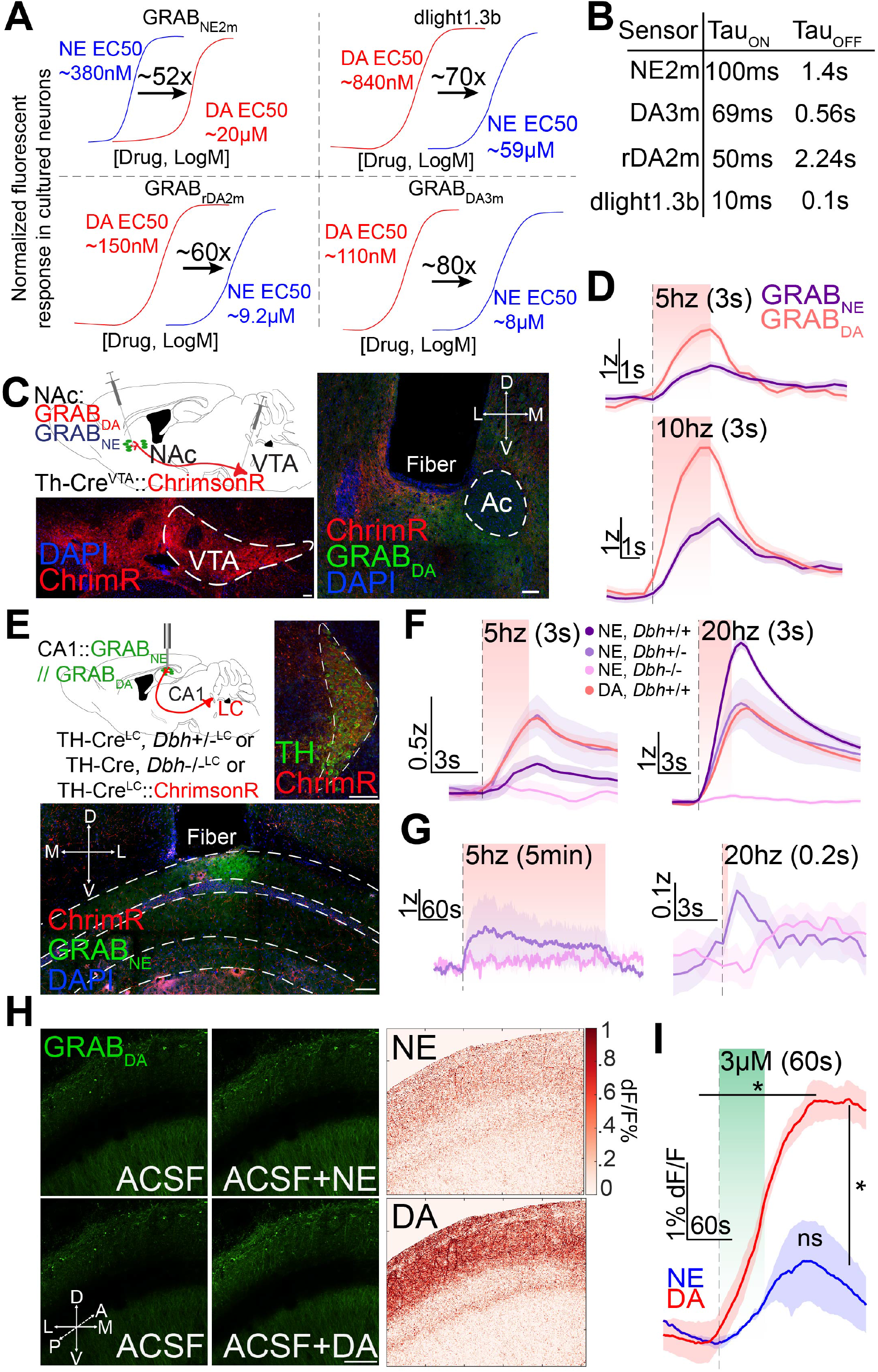
Biosensors are sufficiently selective for their respective ligand. (a) Concentration response curves of various sensors used in this study for NE and DA. EC50s for both NE and DA and the fold differences are recorded. (b) On and off kinetics of the sensors. (c) Schematic depicts infection of VTA-DA cells with ChrimsonR (AAV5-hSyn-FLEX-ChrimsonR-tdTomato) and GRAB_NE_ or GRAB_DA_ in NAc. Representative image of ChrimsonR expression in VTA cell bodies (bottom) and terminals in NAc (right), with GRAB_DA_ expression in NAc. Scale bar, 100μm. (d) VTA-NAc^TH^ terminal stimulation at 5hz for 3s (top) or 10hz for 3s (bottom) E. Schematic of experimental approach for Th-cre, Dbh−/−, and control Dbh+/−heterozygous mice. ChrimsonR was injected into LC (right) and GRAB_NE_ in CA1 (bottom). Scale bar, 100μm. F. Averaged GRAB_NE_ and GRAB_DA_ responses to LC-CA1 stimulation during different stimulation paradigms (left to right) 5hz (3s) and 20hz (3s). G. Averaged GRAB_NE_ responses to tonic (left) and phasic (right) LC-CA1 stimulation during different stimulation paradigms 5hz 5min (left) and 20hz 200ms (right). Dbh(−/−) indicates Dbh-KO mice, n=3, while Dbh+/−indicate heterozygous controls, n=2, and Dbh+/+ are Th-cre mice, n=9 for GRAB_NE2m_, n=3 for GRAB_DA3m_. H. Representative images of baseline (left) and 3μM NE (middle top) or 3μM DA (middle bottom) washes, with heatmaps (right) showing change in fluorescence (ΔF/F). Scale bar, 100μm. I. Averaged traces of 3μM NE and 3μM DA washes. n=2.

### GRAB_NE_ does not detect DA *in vivo* in response to biologically relevant photostimulation

In this study, we employed several catecholamine sensors to understand DA release from LC. The rank order of selectivity (from highest to lowest) of DA sensors is GRAB_DA3m_ (80x NE/DA, EC50NE = 8μM), dlight1.3b (70x, EC50NE = 59μM), GRAB_rDA2m_ (60x, EC50NE = 9.2μM) (Feng et al. 2024 2019, Patriarchi et al. 2018). GRAB_NE2m_ has a 52-fold difference between NE and DA, with EC50DA= 20μM. Because GRAB_NE2m_ has the lowest fold-difference in selectivity, we tested GRAB_NE_’s selectivity by stimulating VTA terminals in BLA and CA1 *in vivo*. We injected either GRAB_NE2m_ or GRAB_DA3m_ in the BLA or CA1 and Cre-dependent viral vectors containing the red-shifted excitatory opsin ChrimsonR, AAV5-hSyn-DIO-ChrimsonR-tdTmto, into the VTA of Th-cre mice (Th-cre::VTA:ChrimsonR) (Figure S1A).

To evoke DA release from VTA terminals, we mimicked sustained low tonic activation (5hz for 30s) and brief, phasic activation (20hz for 3s) that would not inherently be rewarding (Millard et al. 2024) to determine if DA release binds to GRAB_NE2m_. First, we stimulated VTA terminals in BLA and found GRAB_NE2m_ signal increased during both stimulation paradigms (Figure S1B). We found that when we stimulated VTA terminals in CA1, there were also subtle increases in GRAB_NE2m_ fluorescence (Figure S1C). To assess these results further, we repeated this experiment in a region with substantially less NE innervation by stimulating VTA terminals in the nucleus accumbens (NAc) (Figure 1C), which we expected to produce a heightened GRAB_DA3m_ signal. We detected increased GRAB_DA3m_ fluorescence, but a modest GRAB_NE2m_ signal was still detected in response to VTA-NAc stimulation (Figure 1D). Considering that the NAc contains a 1:10 ratio of NE to DA (Koob et al. 1975), we recognized the need to detect GRAB_NE2m_ signal in a completely NE-deficient environment.

To do so, we stimulated LC-CA1 terminals in mice globally deficient in DBH, rendering them unable to produce NE. We expressed AAV5-hSyn-DIO-ChrimsonR-tdTmto in the LC of Dbh(+/+) mice (Dbh-cre::LC:ChrimsonR), Dbh(+/−) mice (Th-cre, Dbh(+/−)::LC:ChrimsonR) that have similar NE content to Dbh(+/+) mice, or Dbh(−/−) mice (Th-cre, Dbh(−/−)::LC:ChrimsonR), and GRAB_NE2m_ in CA1 (Figure 1E). We found that Dbh(−/−) mice had no increase in GRAB_NE2m_ fluorescence when photostimulating for 3 seconds at 5hz or 20hz (Figure 1F). In contrast, GRAB_NE2m_ fluorescence increased in response to LC-CA1 photostimulation in both Dbh(+/+) and heterozygous Dbh(+/−) mice (Figure 1F). Similarly, in response to LC-CA1 photostimulation at both 5hz and 20hz, we detected robust increases in GRAB_DA3m_ signal in Dbh(+/+) mice (Figure 1F-G). These data suggest that in response to LC-CA1 stimulation in the absence of NE production, GRAB_NE2m_ does not detect NE and GRAB_DA3m_ is detecting DA.

The qualitative differences in GRAB_NE2m_ signal between Dbh(+/+) and Dbh(−/−) mice is likely due to differences in strain background (C57BI/6J and mixed C57BI/6J;129SvEv, respectively). To simulate physiological LC activity at high tonic or phasic activation mimicking exposure to a novel-stimulus, we optogenetically stimulated LC terminals in CA1 at 5hz for 5min or at 20hz for 200ms while recording GRAB_NE2m_ signal. We observed that in both activation paradigms, stimulating LC terminals in heterozygous Dbh(+/−) controls increased GRAB_NE2m_ fluorescence, while the same experiment in Dbh(−/−) mice revealed no activation of GRAB_NE2m_ (Figure 1H). These results indicate that the least selective catecholamine biosensor, GRAB_NE2m_ (Figure 1A), does not detect appreciable increases in GRAB_DA3m_ fluorescence at the upper boundary of stimulation parameters where we detect DA release from LC axon terminals (Figure 1F).

To further characterize the selectivity of sensors, we used *ex vivo* slice 2-photon imaging to record sensor activity while washing on high concentrations of ligands. We injected either GRAB_NE2m_ or GRAB_DA3m_ in the CA1 of WT mice. After waiting a minimum 5 weeks for expression, we recorded GRAB_NE2m_ or GRAB_DA3m_ fluorescence in coronal hippocampal slices in response to washes using 2-photon imaging (Figure 1H). When we washed 3μM of NE or DA on slices expressing GRAB_DA3m_, respectively, we observed robust increases in GRAB_DA3m_ fluorescence to DA (Sidak’s multiple comparisons test, p=0.0115) compared to baseline, no significant change in fluorescence from baseline when NE was washed on (p=0.23), and a significant difference between NE and DA (Sidak’s multiple comparisons test, 0.0325). These data suggest GRAB_DA3m_ is responding to DA and not NE in tissue slices at as high a concentration as 3μM.

### *Ex vivo* photostimulation of LC terminals causes release of DA in a frequency-dependent manner

Having demonstrated GRAB_NE2m_ and GRAB_DA3m_’s selectivity over the potential cross-reactive catecholamine, we next sought to better describe the DA release evoked from the LC. To characterize LC-evoked DA release, we used *ex vivo* slice imaging which severs upstream circuit interactions while maintaining local, intact DA release from LC-CA1^Dbh^ terminals. Imaging *ex vivo* also cuts off potential antidromic activation of LC cell bodies and subsequent VTA-DA neuron activation downstream. We injected either GRAB_NE2m_ or GRAB_DA3m_ in the CA1 and Cre-dependent viral vectors containing ChrimsonR, AAV5-hSyn-DIO-ChrimsonR-tdTmto, into the LC of Dbh-cre mice (Dbh-cre::LC:ChrimsonR). After waiting a minimum 5 weeks for expression, we recorded GRAB_NE2m_ or GRAB_DA3m_ fluorescence in coronal hippocampal slices in response to photostimulation of Dbh-cre::LC:ChrimsonR projections (615nm, 5 or 20hz, 5ms pulse width) using 2-photon imaging (Figure 2A). First, we established sensor expression with a positive control wash of 30μM NE or DA on slices expressing GRAB_NE2m_ or GRAB_DA3m_, respectively (Figure 2B). After establishing consistent sensor responses to either NE or DA, we photostimulated LC-CA1^Dbh^ terminals over the full-field, pulsing a 615nm red LED at 5 or 20hz (3 or 30s duration, 5ms pulse width, 3mW) to determine evoked DA release across a range of LC activation intensities. In Figure 2C, an averaged GRAB_DA3m_ trace during 20hz stimulation of LC-CA1^Dbh^ terminals for 30s is provided, indicating robust GRAB_DA3m_ responses to LC-CA1^Dbh^ terminal stimulation.

**FIGURE 2.**
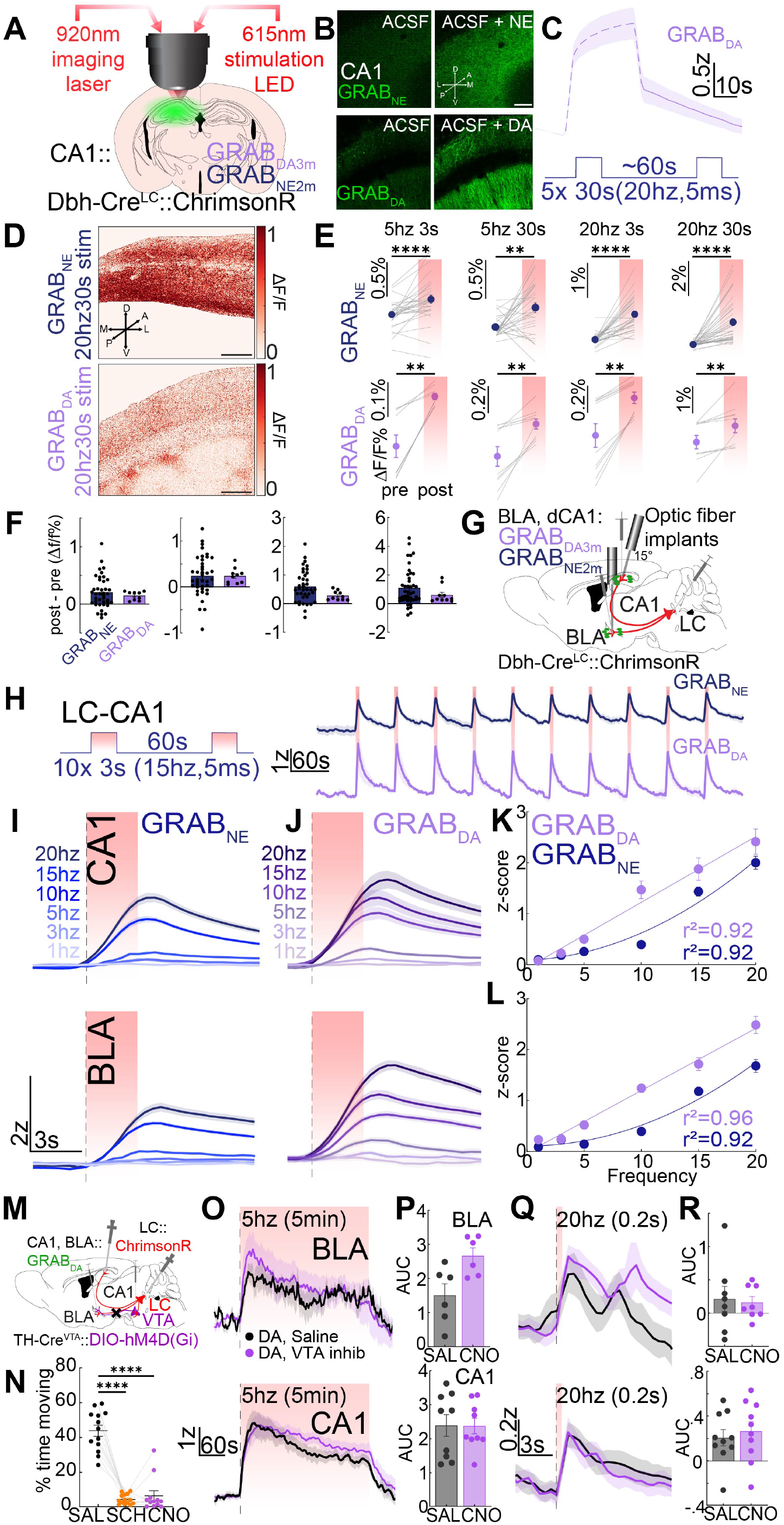
Evoked LC-DA axonal release *in vivo* follows a frequency-dependent linear relationship and is independent of VTA activity. (a) Schematic of experimental approach of surgical injections and 2-photon imaging approach with optical stimulation. Schematic depicts infection of LC-NE cells with ChrimsonR (AAV5-hSyn-FLEX-ChrimsonR-tdTomato) and either GRAB_DA3m_ or GRAB_NE2m_ in CA1. (b) Representative 2-p images. 30μM NE was washed on GRAB_NE_ (top), and 30μM DA on GRAB_DA_ (bottom), expressing slices. Rolling average of 5s, scale bar 100μm. (c) Representative average of GRAB_DA_ response to 20hz 30s pulses of optical stimulation of LC-CA1 terminals. Stimulation paradigm depicted below example trace. (d) Heatmap of the GRAB_NE_ (top) and GRAB_DA_ (bottom). ΔF/F response to 20hz 30s stim of LC-CA1 terminals. Δ represents change from right before stimulation, gaussian filter applied to smooth baseline images. (e) ΔF/F% change of pre to post in GRAB_NE_ (top, n=4 biological replicates, 6 slices) and GRAB_DA_ (bottom, n=2) in response to a range of stimulation paradigms: 5hz 3s, 5hz 30s, 20hz 3s, 20hz 30s (left to right). Wilcoxon matched-pairs signed rank test: GRAB_NE_: 5hz 3s, p<0.0001; 5hz 30s, p=0.0019; 20hz 3s, p<0.0001; 20hz 30s, p<0.0001; GRAB_DA_: 5hz 3s, p=0.0039; 5hz 30s, p=0.002; 20hz 3s, p=0.002; 20hz 30s, p=0.002. (f) Schematic of surgical approach for *in vivo* photometry cohort. GRAB_DA_ or GRAB_NE_ were expressed in CA1 and BLA with fiber photometry implants, and ChrimsonR expressed in LC. Scale bar, 100μm. (g) Comparison of the change in ΔF/F response for GRAB_NE_ and GRAB_DA_ in each stim trial from before to after stimulation across the same paradigms in 2E. Mann-Whitney: 5hz 3s, n.s., p=0.96; 5hz 30s, n.s., p=0.91; 20hz 3s, n.s., p=0.09; 20hz 30s, n.s., p=0.27. (h) Average photometry trace of LC-CA1 stimulation for 3s at 10hz, with a 5ms pulse width. n=5. (i) Averaged *in vivo* photometry traces of GRAB_NE_ responses to LC-CA1 (top) and LC-BLA (bottom) terminal stimulation. (j) Averaged photometry traces of GRAB_DA_ responses to LC-CA1 (top) and LC-BLA (bottom) terminal stimulation. (k) Frequency response curves for LC-CA1 terminal stimulation for GRAB_NE_ and GRAB_DA_. Second-order curve fit to GRAB_NE_, n=5-9, r^2^= 0.92, and linear regression fit to GRAB_DA_, n=3, r^2^= 0.92. (l) Frequency response curves for LC-BLA terminal stimulation for GRAB_NE_, n=6, and GRAB_DA_, n=3. Second-order curve fit to GRAB_NE_, r^2^= 0.92, and linear regression fit to GRAB_DA_, r^2^= 0.96. (m) Schematic of experimental approach of surgical injections of LC-NE cells with ChrimsonR (AAV5-hSyn-FLEX-ChrimsonR-tdTomato), VTA-DA cells with Gi-DREADDs (AAV5-hSyn1-DIO-HA-hM4D(Gi)), and GRAB_DA3m_ in CA1, of Th-cre mice. (n) VTA inhibition decreased the time mice spent moving. SAL, n=7; SCH, n=7; CNO, n=6. (o) Average trace of GRAB_DA_ response to tonic stimulation (5hz, 5min) of LC terminals in BLA (top; n=8) and CA1 (bottom, n=10), with VTA inhibition (purple) or saline controls (black). (p) AUC for 120s after start of 5hz stimulation for mice injected with saline or CNO (VTA Inhibition). (q) Average trace of GRAB_DA_ response to phasic stimulation (20hz 200ms) of LC terminals in BLA (top; n=8), and CA1 (bottom, n=10). (r) AUC for 3s after start of 20hz, 200ms stimulation for mice injected with saline or CNO (VTA Inhibition).

In both GRAB_DA3m_ and GRAB_NE2m_ expressing CA1 slices, we observed a significant increase in fluorescence across all stimulation paradigms tested (Figure 2D-E). We compared the change in ΔF/F% at each of the stimulation paradigms and found that none of the ΔF/F% changes between GRAB_NE2m_ and GRAB_DA3m_ response were significantly different (Figure 2F). To test the consistency of these results, we repeated these *ex vivo* experiments with dLight expressed in CA1, and ChrimsonR expressed in the LC of Dbh-cre mice (Figure S2A-B, I). We only observed significant increases in fluorescence when we stimulated for 3s at 20hz, while other paradigms were more variable in their detected DA release (Figure S2C). These results indicate DA is released from LC-CA1^Dbh^ terminals *ex vivo*, regardless of any connections from LC to other dopaminergic nuclei.

### Evoked LC-NE axonal release *in vivo* follows a frequency-dependent, non-linear relationship

Our *ex vivo* results demonstrate that photostimulating LC-CA1^Dbh^ axon terminals evokes DA release. We next determined how LC-evoked DA release varies across a range of photostimulation paradigms that parallel natural neuronal LC activity *in vivo* (Bourdélat-Parks et al. 2005, Devilbiss and Waterhouse 2011, Szot et al. 1999, Thomas et al. 1998 1995) and examined two target regions. We injected either GRAB_NE2m_ or GRAB_DA3m_ in CA1 or BLA, and Cre-dependent viral vectors containing ChrimsonR, into the LC of Dbh-cre mice (Dbh-cre::LC:ChrimsonR; Figure 2G). We implanted optical fibers above injection sites in BLA and CA1 to record photometric signals during optogenetic stimulation of LC axon terminals.

During periods of low arousal, such as sleep, LC fires tonically at 1-3hz. In high arousal states, like stress, elevated tonic LC-NE neuron firing rates are observed (between 5-10hz) (Aston-Jones and Cohen 2005, Curtis et al. 2012, Devilbiss and Waterhouse 2011, Poe et al. 2020). Recent studies report release of DA from LC-CA1 projections in response to photostimulation at 20hz (Kempadoo et al. 2016, Takeuchi et al. 2016). Thus, we stimulated the LC-CA1^Dbh^ and LC-BLA^Dbh^ projections for 3s at 1, 3, 5, 10, 15, and 20hz with a 5ms pulse width and a total of 10 stimulation trains each session with an interstimulation interval of 1min (Figure 2H-L). We depict a representative averaged response of GRAB_NE2m_ and GRAB_DA3m_ to the stimulation at 15hz for 3s in Figure 2H. We also depict the responses to LC-CA1 stimulation averaged across frequency in Figure 2I for GRAB_NE2m_ recordings and Figure 2J for GRAB_DA3m_ recordings. To characterize trends within each sensor and across regions, we plotted the peak responses across the frequencies tested in CA1 (Figure 2K) and in BLA (Figure 2L). LC-BLA^NE^ and LC-CA1^NE^ release across frequencies strongly matched a second-order polynomial fit (adjusted R^2^ values in CA1 was 0.92, BLA was 0.92). LC-CA1^DA^ and LC-BLA^DA^ release across frequencies more closely followed a linear regression model, with adjusted R^2^ values of 0.92 in CA1 and 0.96 in BLA (Figure 2L). Further, optically evoked LC-CA1^DA^ release *in vivo* was not significantly different from evoked LC-BLA^DA^ release (Kolmogorov-Smirnov test, n.s., p=0.93; Figure 2K-L). These data suggest that the distributions of LC evoked DA release in different regions are not significantly different.

We repeated these experiments in mice expressing dLight in CA1 and BLA, with Cre-dependent ChrimsonR expressed in the LC of Dbh-cre mice (Figure S2D, I). We repeated *in vivo* stimulation for 3s across the same range of frequencies reported in Figure 2H-L (1-20hz) and depicted averaged responses in Figure S2E, G. We characterized trends in DA dynamics response across frequency and found that, a second-order polynomial was not appropriately fit in the BLA (adjusted R^2^ values in BLA was 0.30), although in CA1, a second-order fit was moderately well-fit (R^2^ values in CA1 was 0.84; Figure S2F, S2H). These data suggest that while NE release from LC follows a second-order relationship, LC evoked DA release follows a linear relationship. This points to likely differences in uptake, autoreceptor activity, metabolism, repletion, or sensor kinetics; the findings nevertheless indicate a potential separation of relevant LC firing properties and the relative NE and DA concentration in each. Further studies using the newest high-resolution FLIM based sensors are needed to mechanistically quantify these findings (Lee et al. 2019, Salinas et al. 2023, Zheng et al. 2025).

### DA release from LC axons persists during VTA inhibition

In the 2-photon slice experiments (Figure 2D-H), we severed connections from LC to CA1, preventing antidromic activation of the LC, but local DA neuron axon terminals may still be activated by interactions with evoked NE release via 2AR or 1AR expression on VTA neurons (Bernacka et al. 2022, Boyajian and Loughlin 1987, Greene et al. 2005, Nicholas et al. 1993ba, Rosin et al. 1993). To remove VTA as a source of DA when LC terminals are stimulated, we inhibited the VTA while recording GRAB_DA_ signal. We expressed the Cre-dependent inhibitory designer receptor exclusively activated by designer drugs (DREADD), virally delivered with AAV5-hSyn-DIO-HA-hM4D(Gi) bilaterally in the VTA of Th-cre mice (Th-cre::VTA^bilat^:Gi-DREADDs), AAV5-hSyn-DIO-ChrimsonR-tdTmto in the LC (Th-cre::LC:Chrimson), and AAV2/9-hSyn-WPRE-PA-GRAB_DA3m_ in CA1 or BLA (Th-cre::CA1 or BLA: GRAB_DA3m_; Figure 2M). When we injected the DREADD agonist clozapine-N-oxide (CNO) (0.5 mg/kg, i.p.) 30min prior to exposing mice to a large open field chamber (50×50cm), we observed a significant decrease in the time the mice spent moving, which were mimicked by the effects of the D1-antagonist SCH23390 (0.25 mg/kg, i.p.) (Figure 2N). D1 dopamine receptors are involved in movement initiation and execution (Hauber 1996, Heath et al. 2015), thus these data indicate that the hM4Di expressed is reliably suppressing DA transmission.

We next tested whether VTA-DA neurons were required for LC-evoked DA release and determined if VTA-DA neuronal involvement in LC-evoked DA release depends on the mode of LC activity. To induce a range of LC activity, we simulated low tonic, high tonic, and phasic stimulation. For low tonic or high tonic stimulation, we stimulated terminals for 5min at either 1hz or 5hz. Photostimulation of LC-CA1^Th^ and LC-BLA^Th^ terminals evoked DA release in response to 5hz (Figure 2O-P). To simulate phasic stimulation, we stimulated LC terminals in BLA and CA1 for 200ms at 20hz, which evoked DA release in both (Figure 2Q-R). When we inhibited the VTA using CNO, evoked DA did not decrease in either 5hz tonic stimulation (Figure 2O-P) or in 20hz phasic stimulation (Figure 2Q-R), indicating the VTA is not necessary for evoking DA release from LC axons. Taken all together, these results further establish that the LC releases DA independent of VTA activity.

### Appetitive and aversive stimuli evoke more DA release in BLA compared to CA1

Previously, LC-evoked DA has been suggested to affect behavior, but there are no data that report endogenous DA release during behavior (Kempadoo et al. 2016, Takeuchi et al. 2016). To ascertain whether the LC releases DA endogenously in response to salient stimuli, we first investigated endogenous release of DA and NE in BLA and CA1 in response to simple aversive and appetitive stimuli. To detect NE or DA dynamics, we injected virally-delivered green NE or DA sensors, AAV1-hSyn-GRAB_NE2m_ or AAV2/9-hSyn-WPRE-PA-GRAB_DA3m_, respectively, into the BLA or CA1 (Dbh-cre::CA1 or BLA:GRAB_NE2m_ or GRAB_DA3m_; Figure 3A) of Dbh-cre mice we used in photostimulation experiments in Figure 2. We implanted optical fibers above injection sites in BLA and CA1 to record photometric signal during stimulus administration, including footshock, reward (Figure 3B).

**FIGURE 3.**
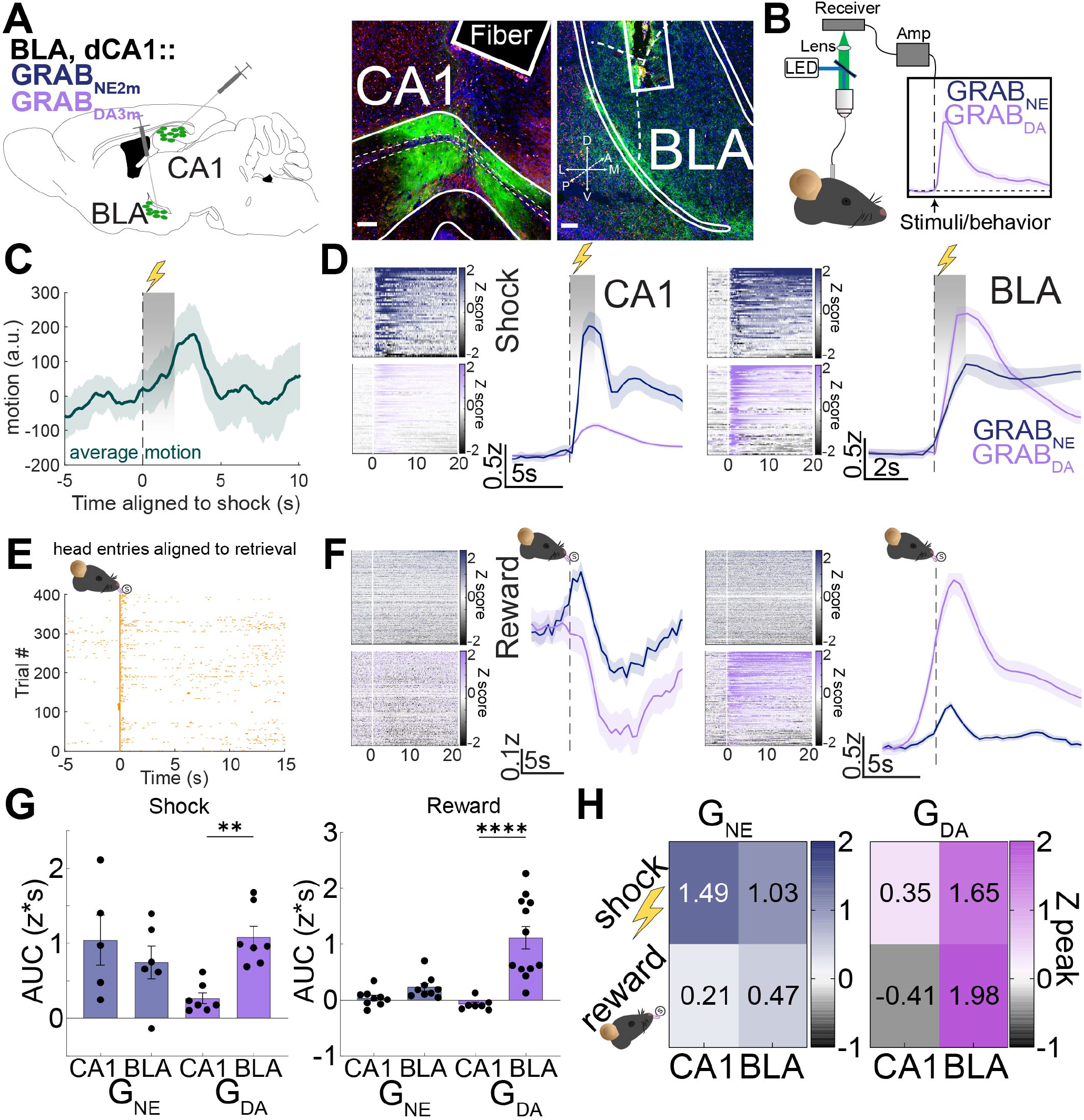
Reward and aversive stimuli evoke primarily NE in CA1 and DA in BLA. (a) Schematic of experimental approach depicts infection of cells with GRAB_DA_ (AAV2/9-hSyn-WPRE-PA-DA3m) or GRAB_NE_ (AAV1-hsyn-GRAB_NE2m_) in CA1 and BLA with fiber photometry implants. Scale bar, 100μm. (b) Schematic of photometry acquisition. (c) Average motion aligned to shock in mice experessing GRAB_DA_. (d) GRAB_NE_ and GRAB_DA_ traces in response to 2s shock (0.5mA to 0.7mA) presented with a pseudorandom ITI of 20-90s in CA1 (left, nNE=5, nDA=7) and BLA (right, n_NE_=6, n_DA_=7). Heatmaps for GRAB_NE_ and GRAB_DA_ responses to shock with color axes from −2 to 2. (e) Example rasterplot of reward delivery and subsequent head entries in trained mouse. (f) GRAB_NE_ and GRAB_DA_ response to reward in CA1 (left, n_NE_=9, n_DA_=7) and BLA (right, n_NE_=9, n_DA_=12). Heatmaps for GRAB_NE_ and GRAB_DA_ responses to reward retrieval with color axes from −2 to 2. Sucrose pellets were delivered to hungry mice with a pseudorandom ITI of 30, 60, 90, 120s with an average of 90s. (g) Quantification of the AUC of the average GRAB_NE_ or GRAB_DA_ response to shock (left) or reward (right). Post hoc P-values derived from Kruskal-Wallis with Dunn’s multiple comparison. Shock, GRAB_DA_-CA1 vs -BLA, p=0.0095. Reward, GRAB_DA_-CA1 vs GRAB_DA_-BLA, p<0.0001. (h) Heatmap of peak z-scores during both experiments in 3D and 3F. All data are mean ± SEM.; n represents biologically independent mice.

First, mice were exposed to a 2s footshock, a highly salient and aversive stimulus. During the 2s shock, mice moved more than before each shock (Figure 3C). In CA1, GRAB_NE2m_ and GRAB_DA3m_ fluorescence increased at the onset of the shock, with the GRAB_DA3m_ response lower than GRAB_NE2m_ (Figure 3D, left, and Figure S3A-B). In BLA, the GRAB_DA3m_ response was greater than GRAB_NE2m_ in response to shock, but GRAB_NE2m_ had a slow decay for over 30s, which was notably different to the decay in GRAB_NE2m_ signal in response to other stimuli (Figure 3D, right, and Figure S3C-D). Only GRAB_NE2m_ had a sustained and significantly increased signal for the rest of the session compared to baseline (Figure S3I). In response to shock, the BLA-DA response was higher than the CA1-DA response (Figure 3G, Kruskal-Wallis with Dunn’s multiple comparison, p=0.0095). These data indicate NE and DA are released in both CA1 and BLA in response to aversive stimuli.

Next, we investigated baseline NE and DA dynamics in response to an appetitive stimulus, the retrieval and consumption of a sucrose reward. We recorded NE and DA dynamics as mice completed the task, and aligned photometry signals to the moment mice entered their heads in the hopper to retrieve sucrose pellets throughout the session (Figure 3E). We found that as mice approached the hopper for reward, NE was released in CA1, and as mice consumed, there was a subtle dip in the GRAB_NE2m_ signal (Figure 3F, left). In the BLA, both NE and DA increased as mice approached the hopper to retrieve a sucrose pellet (Figure 3F, right). In response to shock and reward, GRAB_DA3m_ signal increased in BLA greater than in CA1 (Figure 3G, shock: p=0.0095, reward: p<0.0001), whereas GRAB_NE2m_ signal was not significantly different within each behavior across regions. These data suggest more DA is released in BLA than in CA1 in response to salient stimuli (Figure 3G-H).

In addition to shock and reward delivery, we tested additional types of appetitive and aversive behaviors. First, we used a looming stimulus to simulate a less salient, aversive stimuli, and found that NE was released in CA1 and BLA to the onset of the looming stimulus (Figure S3E). Similar to the footshock, GRAB_DA3m_ signal increased in BLA, but not in CA1, at the onset and offset of the looming stimulus (Figure S3E). Mice also exhibited darting behavior at the onset of the looming stimuli (Figure S3F). To assess a second appetitive behavior, mice were trained to poke their nose in a port for a reward predicted by a 2s light cue, while a second, inactive nose port, led to no reward delivery. When trained mice poked their nose in the correct port, dLight signal in BLA increased in anticipation of the cue-signaling reward (Figure S3G). While the dLight and GRAB_NE_ signal in BLA did not increase to cue, the dLight and GRAB_NE_ signals in BLA increased to reward retrieval and consumption (Figure S3G, top). There was no increase in dLight signal in CA1 during operant paradigm, but there was a decrease in GRAB_NE_ signal to nosepoking for reward (Figure S3G, bottom). When mice poked their noses in the inactive nose port, there was a clear increase in dLight in BLA and a smaller increase in CA1, while there was no increase in GRAB_NE_ signal in BLA or CA1 (Figure S3H). All together, these results establish that we can detect DA that is released in the BLA following both aversive and appetitive stimuli, and DA is only detected in CA1 during salient aversive stimuli.

### LC inhibition decreases BLA DA release to appetitive and aversive stimuli

Finally, we sought to conclusively demonstrate that DA released following appetitive and aversive stimulus presentation is from LC-BLA axon terminals. To accomplish this, we used inhibitory chemogenetics to suppress either LC-NE or VTA-DA activity. To accomplish this, we injected the Cre-dependent Gi-DREADD (AAV5-hSyn1-DIO-HA-hM4D(Gi)) bilaterally in the LC of Th-cre mice (Th-cre::LC^bilat^:Gi-DREADDs) (Figure 4A) and AAV9-hSyn-GRAB_rDA2m_ in the CA1 (Figure 4B) or BLA (Figure 4C).

**FIGURE 4.**
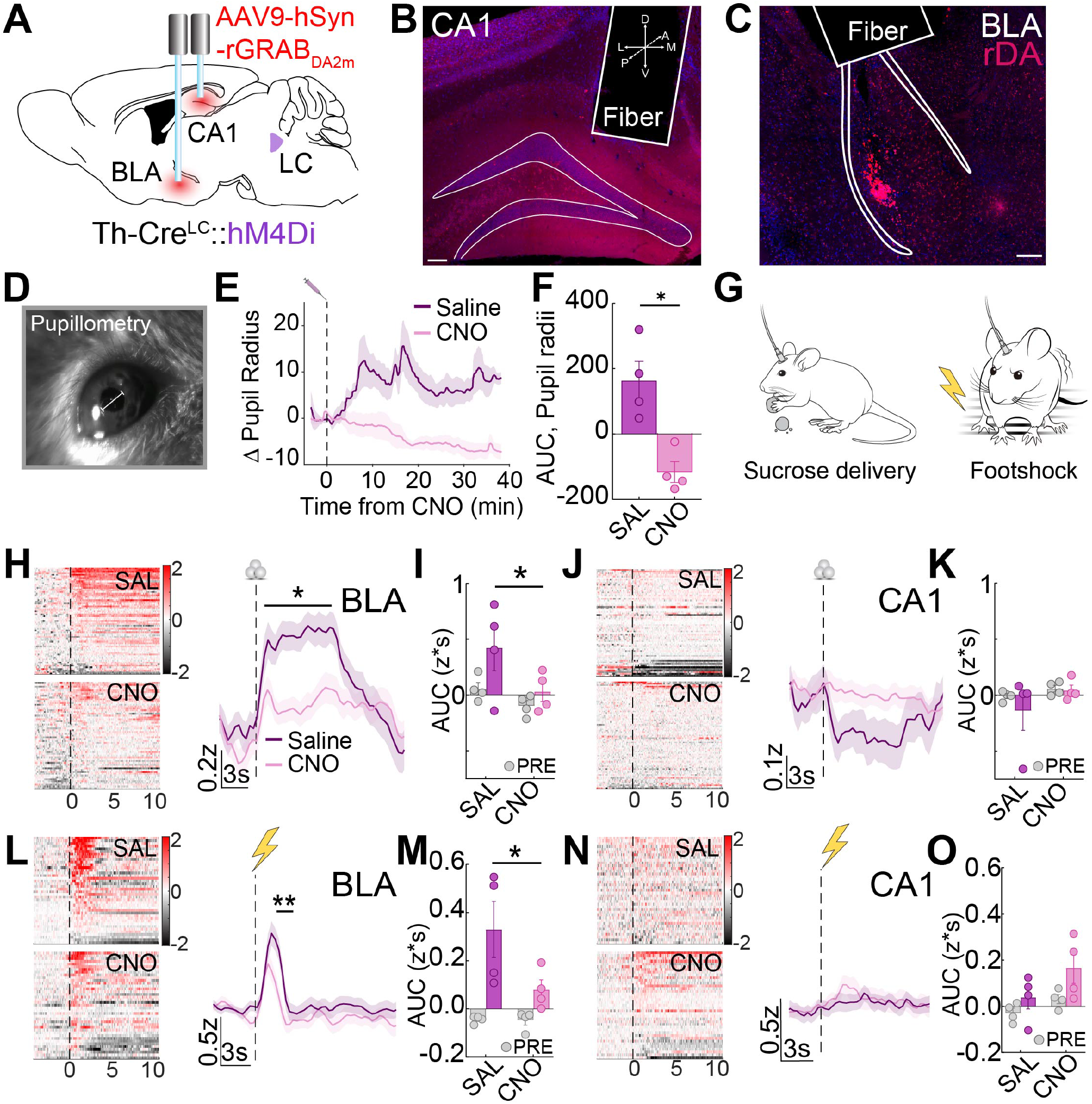
LC inhibition decreases DA release during reward and shock in BLA, and has no effect in CA1. (a) Schematic of experimental approach of surgical injections. Schematic depicts infection of LC-NE cells with Gi-DREADDs (AAV5-hSyn1-DIO-HA-hM4D(Gi)) and expression of GRAB_rDA2m_ in both BLA and CA1. (b) Representative image of CA1 GRAB_rDA2m_ expression and fiber placement. Scale bar, 100μm. (c) Representative image of BLA GRAB_rDA2m_ expression and fiber placement. (d) Example of pupillometry recording with measurement depicted. (e) Pupil radius measurements during LC inhibition. Gi-DREADDs were expressed bilaterally in LC. n=4. (f) AUC for pupil radius measurements. (paired t-test, p=0.0223). (g) Schematic of sucrose delivery (H-K) and footshock (L-O). (h) Average traces and heatmaps for GRAB_rDA2m_ response to reward retrieval in BLA during LC inhibition (pink) and saline controls (purple). Significance of 1s bins calculated using 2-way ANOVA, with Sidak test for multiple comparisons, p<0.05. n=4. (i) AUC for 10s in SAL and CNO responses to reward in BLA. 2-way ANOVA, with Sidak test for multiple comparisons, p=0.0327. (j) Average traces and heatmaps for GRAB_rDA2m_ response to reward retrieval in CA1 during LC inhibition or saline. n=4. (k) AUC for 10s in SAL and CNO responses to reward in CA1. (l) Average traces and heatmaps for GRAB_rDA2m_ response to shock in BLA during LC inhibition (pink) and saline (purple). Significance of 1s bins calculated using 2-way ANOVA, with Sidak test for multiple comparisons, p<0.05. n=4. (m) AUC for 5s in SAL and CNO responses to reward in BLA. 2-way ANOVA, with Sidak test for multiple comparisons, p=0.025. (n) Average traces and heatmaps for GRAB_rDA2m_ response to shock in CA1 during LC inhibition or saline. n=4. (o) AUC for 5s in SAL and CNO responses to reward in CA1.

Because pupil dilation is positively correlated with higher LC activity(Liu et al., 2017; Murphy et al., 2014), we used pupillometry to establish the efficacy of Gi-DREADD inhibition of LC activity (Privitera et al. 2020) (Figure 4D). When we injected the DREADD agonist CNO (0.5 mg/kg, i.p.) in Th-cre mice expressing hM4D(Gi) in LC bilaterally, we observed a consistent, significant decrease in pupil radii compared to saline, as measured by comparing the “area under the curve” (ΔAUC, z*s), suggesting that the LC was inhibited by this chemogenetic approach (Figure 4E-F, paired t-test, p=0.0223).

With DREADD suppression of the LC-NE system established, we next injected either CNO (0.5 mg/kg, i.p.) or saline 30 min prior to a session in which mice received pseudorandom (ITI of 90-150s) delivery of sucrose pellets while recording GRAB_rDA2m_ (Figure 4I) to measure BLA DA release during reward consumption. Following saline administration, as mice retrieved a reward and throughout consumption, DA was released in the BLA before returning to baseline (Figure 4H). Relative to saline treatment, we observed a significant decrease in BLA DA release when mice retrieved and consumed a reward following CNO inhibition of LC (Figure 4H, 1s bins analyzed with 2-way ANOVA, Sidak test for multiple comparisons, p<0.05 for all bins). Comparing the total change in DA release (ΔAUC, z*s), we found LC inhibition significantly reduced DA release in BLA during reward consumption relative to saline (Figure 4I, 2-way ANOVA, with Sidak test for multiple comparisons, p=0.0327). In CA1, we did not observe DA release during reward consumption (Figure 4J-K). In summary, these data indicate that inhibiting the LC during reward retrieval leads to a decrease in DA release in BLA and no change in CA1.

VTA activity and dopamine release is correlated with salient stimuli (Kutlu et al. 2021) and lesioning the LC or VTA disinhibits the activity of the other (Guiard et al. 2008). Thus, VTA activity’s influence on DA release to relevant stimuli is important to understanding the LC’s role in DA release. To determine the influence of VTA on DA release in response to reward, we inhibited the VTA of Th-cre mice (Th-cre::VTA^bilat^:Gi-DREADDs) as mice retrieved rewards and recorded GRAB_rDA2m_ dynamics (Figure S4A). Following saline administration, DA was released in BLA during reward retrieval, and the GRAB_rDA2m_ signal remained elevated during consumption (Figure S4B, left). In CA1, DA activity did not change (Figure S4B, right). In response to VTA inhibition, there was a decrease in DA that was released in the BLA during reward consumption compared to saline administration sessions, while there was no effect of this manipulation on DA activity in CA1 (Figure S4B). The latency of mice to retrieve reward was significantly increased when the VTA was inhibited (Figure S4C), establishing that the VTA’s activity was suppressed by DREADD manipulation. Together, these data indicate that the LC, with the VTA, both contribute DA in the BLA during reward retrieval.

To test whether the GRAB_rDA2m_ sensor is detecting NE under the same conditions, we recorded GRAB_rDA2m_ and GRAB_NE2m_ signal in the BLA simultaneously while mice retrieved a pellet for reward using dual-color photometry recordings. We found a clear separation in the onset of the sensors’ signals (Figure S4D) that was greater than the reported on-times of the sensors (Figure 1B), suggesting these signals are likely separable and independent if carefully monitored using these considerations. Furthermore, as mice consumed reward, GRAB_rDA2m_ sensor activity remained elevated for 20s after the GRAB_NE2m_ activity returned to baseline. Lastly, there was no correlation between the AUC (10s post reward retrieval) of the simultaneously recorded NE and DA signals (Figure S4E). These recordings indicate we can detect separable GRAB_NE2m_ and GRAB_rDA2m_ signals simultaneously during freely moving behaviors, assuming sensor kinetics and transmitter release properties are divergent.

Next, we investigated the VTA’s putative involvement in DA release to appetitive and aversive stimuli. Prior studies implicated DA release from LC-CA1 axons in spatial memory without VTA involvement (Kempadoo et al. 2016, Takeuchi et al. 2016), while other studies have suggested DA sourced from the VTA axon terminals modulate aversive memory acquisition independent of the LC (Tsetsenis et al. 2021). To test whether LC-sourced DA is released in response to aversive stimuli, we again used chemogenetic inhibition of LC (Th-cre::LC^bilat^:Gi-DREADDs) prior to the delivery of an aversive footshock stimulus while recording GRAB_rDA2m_ dynamics in CA1 and BLA. CNO (0.5 mg/kg, i.p.) or saline were injected 30min prior to each session, and mice received footshocks at a pseudorandom ITI of 20-90s. Following saline administration, DA was released in the BLA at the onset of a footshock, and the DA signal fell rapidly following the shock delivery. In the CA1, DA release was surprisingly not detected under the same conditions during footshock. Relative to saline treatment, inhibiting the LC-NE system during shocks produced a significant decrease in DA release in BLA during the two seconds from shock initiation (comparing 1s bins, Figure 4L, 2-way ANOVA, p<0.01). LC inhibition also produced a significant decrease in the GRAB_rDA2m_ signal in BLA for 5s after shock onset compared to saline (Figure 4M, 2-way ANOVA, p<0.05). In CA1, LC inhibition had no effect on the release of DA (Figure 4N-O).

Finally, we next measured how VTA inhibition affected DA release in response to an aversive stimulus. In the BLA, DA increased with the onset of shock and returned to baseline after cessation, while in the CA1, GRAB_rDA2m_ signal did not change with the shock. When the VTA was inhibited during shock sessions, there were no clear changes in the GRAB_rDA2m_ signal detected in BLA or CA1 relative to saline (Figure S4F). These results indicate that VTA inhibition does not affect DA release in response to shock in both BLA and CA1 compared to saline controls, while during reward retrieval and consumption, VTA inhibition decreased DA release in the BLA compared to saline trials. Thus, only LC inhibition decreases DA release in the BLA following an aversive stimulus, suggesting that LC-NE neurons represent a major source of local DA release within the BLA during aversive stimuli.

## 3 DISCUSSION

In summary, here we used optogenetics, fiber photometry and 2-photon *ex vivo* imaging to discover that stimulation and natural behavior both elicit dynamic DA release from LC terminals. We demonstrate that *in vivo*, LC-evoked NE release followed a second-order frequency response curve, while the LC-evoked DA release followed a linear trend in both BLA and CA1. We also established that LC inhibition diminished DA release in response to reward retrieval and shock in the BLA, while having minimal influence on DA release in the CA1. Together, we have established here that the LC has the capacity to release DA in both CA1 and BLA under physiologically meaningful conditions. We also determined that during appetitive and aversive stimuli, LC contributes DA release in BLA, but not appreciably in the CA1.

Recent studies have asserted that LC-NE neurons release DA in the hippocampus, enhancing contextual memory and plasticity independent of the VTA (Kempadoo et al. 2016, Smith and Greene 2012, Takeuchi et al. 2016). Some studies have used optogenetics and chemogenetics to evoke DA release from LC in dorsal CA1, but only over very long temporal domains (Kempadoo et al. 2016, Wilmot et al. 2024). In the present study we optically stimulated brain slices at much lower stimulation duration (3s), more indicative of LC-NE neuron burst activity and frequency (as low as 1hz) to determine the lower limits of physiologically-relevant LC-evoked DA and NE release. This study further characterizes LC-evoked DA release by mimicking phasic (20hz, 200ms) and tonic (5hz, 5min) activation profiles matched to physiologically relevant parameters of previously reported LC activity across mammalian species (Figure 1F, 2O-R). Although we acknowledge that optogenetic stimulation is inherently artificial, we used paradigms in this study which fall within the range of previously used and well-characterized optogenetic stimulation paradigms for LC (Carter et al. 2010, McCall 2015) actuation, which we carefully match to the natural range of LC activity (Carter et al. 2010, Devilbiss and Waterhouse 2011). In response to stimulation mimicking the natural range of LC activity, the evoked LC-NE signal followed a second-order trend while the evoked LC-DA signal followed a linear trend. These data suggest that NE and DA release from LC terminals are regulated differently, and that there are likely differences in uptake, autoreceptor activity, metabolism, or repletion kinetics (as we recently reported) (Li et al. 2023). Future studies will need to characterize the differences in NE and DA release during various stimulation paradigms while altering transporter and vesicle activity, and also utilize FLIM-based sensor technologies, to accurately quantify these differences with higher resolution (Lee et al. 2019, Salinas et al. 2023, Zheng et al. 2025).

Additionally, in situ RNA hybridization, autoradiography and immunohistochemistry have revealed varying degrees of α1AR expression on VTA neurons (Boyajian and Loughlin 1987, Greene et al. 2005, Mitrano et al. 2012, Nicholas et al. 1993ba, Rosin et al. 1993, Solecki et al. 2022). The expression of adrenergic receptors on VTA terminals suggests LC stimulation may amplify DA release from VTA-DA terminals via a Gq-mediated effect. However, by inhibiting VTA^TH^ neurons while stimulating LC terminals, we determined that LC-evoked DA release in CA1 and BLA does not entirely rely on VTA activation (Figure 2O-2R).

While receptor pharmacology experiments have revealed the potential for DA release from noradrenergic terminals, electrical stimulation, lesions of LC, and genetic knockout of Dbh have been used to demonstrate the necessity of the LC for DA release. Selective lesioning of the LC leads to an increase in VTA-DA neuron activity, and VTA-DA lesions lead to an increase in LC-NE neuron activity (Grenhoff et al. 1993), suggesting that there are functional LC-VTA reciprocal projections (Deutch et al. 1986, Mejias-Aponte et al. 2009). These data would predict that in response to a stimulus that evokes VTA-BLA^DA^ release, the VTA may release more DA when the LC is inhibited. However, our data suggest that either LC or VTA inhibition decrease the BLA-DA release during reward retrieval (Figure 4H-I, Figure S4C). Furthermore, only LC inhibition decreased the BLA-DA release during shock (Figure 4L-M, Figure S4D), suggesting the LC’s contribution of DA during an aversive stimulus may be greater than the DA released following disinhibition of VTA. Therefore, these findings taken together indicate that the LC releases DA in the BLA in response to both aversive and appetitive stimuli.

Although LC inhibition did not affect CA1-DA release in response to aversive and appetitive stimuli with GRAB_rDA2m_ (Figure 4J, 4N), DA was evoked from LC terminals in CA1 through optogenetic stimulation *ex vivo* and *in vivo*. This suggests that LC-CA1^DA^ release is dependent on behavioral states and neuronal activation. Because stimuli-evoked DA release decreases during LC inhibition in BLA and not CA1, but optically evoked DA release frequency response profiles do not differ significantly in these two regions (Figure 2K, 2L), LC-CA1 projections’ endogenous release of DA may rely on different behavioral states than LC-BLA projections.

Recording a ligand-specific signal relies on silencing cross-reactive neuromodulator sources (e.g. VTA-DA and LC-NE neurons both project to CA1 and BLA) (López et al. 2024). To address these concerns with the biosensors used in this project, we employed genetic knockouts, inhibitory chemogenetics, and dual-color fiber photometry recordings. Through stimulating LC-CA1^Dbh−/−^ projections for 3s at 20hz, there was no increase in NE signal at the upper limit of stimulation frequencies employed. In addition, GRAB_NE_ and GRAB_DA_ dynamics differ in response to reward retrieval and looming in CA1 (Figure 3F, S3E). This suggests that in our stimulation experiments, GRAB_NE_ is sufficiently selective for NE over DA. To address concerns that the red GRAB_rDA2m_ sensor used in Figure 4H-O and Figure S4B, S4D is detecting NE, we established that GRAB_rDA2m_ dynamics are separable from GRAB_NE_, by using dual-color fiber photometry recordings in naïve, hungry mice retrieving rewards (Figure S4C). Given the known off-kinetics of these biosensors (< 3s, Figure 1B) and the period of heightened GRAB_rDA2m_ signal (>20s), GRAB_rDA2m_ is likely not binding significant NE at this level of endogenous release. These data taken together suggest the GRAB_rDA_ and GRAB_NE_ are sufficiently selective for endogenous detection of NE and DA in these experiments.

There are also other sources of DA and NE than the LC and VTA that must also be considered herein. Studies suggest that Dorsal Raphe nucleus (DRN) has either negligible (Cho et al. 2017, Sengupta and Holmes 2019) DA innervation in BLA, and none in CA1. Substantia nigra (SN) projects to BLA, but not CA1, and these BLA projections are non-DAergic projections (Loughlin and Fallon 1983). Further studies are needed to address if the DRN or SN inputs in BLA influence the interpretation of LC-BLA^DA^ release results we report.

Past studies have suggested that DA release in BLA contributes to fear expression, primarily acquisition (Guarraci et al. 2000, Heath et al. 2015, Lamont and Kokkinidis 1998, Nader and LeDoux 1999), and that BLA-projecting VTA neurons, specifically, have been shown to release DA and contribute to conditioned fear acquisition (Tang et al. 2020). Another study demonstrated BLA projecting LC neurons contribute to fear memory formation (Uematsu et al. 2017). Our data suggest the LC is contributing some portion of DA in BLA during an aversive unconditioned stimulus (Figure 4L). Given both NE and DA are involved in fear acquisition in the amygdala (Giustino et al. 2020, Giustino and Maren 2018, Heath et al. 2015, Pantoni et al. 2020), future studies should determine how LC-sourced NE and DA released in the BLA may be involved in fear acquisition. Especially in regions with dual VTA and LC innervation, investigators focused on VTA-DA must carefully consider noradrenergic sources of DA in interpreting results.

Activity of CA1 neurons during aversive stimuli like tail shock and airpuff scales with salience and context independent of location (Barth et al. 2023), increases with the valence of a reward-predictive cue (Yun et al. 2023), and certain populations respond to reward (Gauthier and Tank 2018). While DA has been suggested to be involved in enhancing memory persistence and spatial memory (Kempadoo et al. 2016, Takeuchi et al. 2016), the present study is among the first to look at dopamine’s release dynamics within CA1 during aversive and appetitive stimulus presentation. While DA was not detected in response to reward in CA1, DA release was moderately present during shock in CA1, which may be sourced from the VTA and modulating the formation of aversive memories (Tsetsenis et al. 2021). Together these data indicate that LC-CA1 projections have the capacity to release DA, but only do so under specific conditions. As hippocampal inputs to BLA are important for contextual conditioning, CA1-DA may signal the valence of contextual information, which is then relayed to the BLA for consolidation.

While this study demonstrates the conditions for DA release from LC terminals, we do not address mechanisms of LC-DA release. DBH is not rate-limiting for NE synthesis thus how the LC may release DA has remained controversial even with emerging evidence of DA release from noradrenergic terminals. DBH is primarily localized to be inside of synaptic vesicles (Cimarusti et al. 1979) although it has also been observed less-selectively throughout the somata and processes of LC neurons (Olschowka et al. 1981), while TH is localized in the cytoplasm (Pickel et al. 1975). For DA to be released from LC neurons endogenously as the present study suggests, there must be some population of synaptic vesicles in LC terminals that do not contain DBH or whole terminals that exclude DBH-containing vesicles. Future work should use electron microscopy to explore the distribution of vesicles and DBH, and TH within LC terminals in various regions throughout the brain.

Lastly, the LC contributing DA in BLA during reward suggests there must be some active process by which DA is endogenously released from LC terminals. In some human populations, pharmacological (Gaval-Cruz and Weinshenker 2009) (e.g. disulfiram) and genetic factors (Cubells and Zabetian 2004) (e.g. *Dbh* deficiency, *Dbh* gene polymorphisms) may affect the ratio of DA/NE released by the LC, which could impact behavior especially with maladaptive comorbidities like addiction or ADHD. Due to the LC’s characteristic functional differences at specific projections (Chandler et al. 2019, Uematsu et al. 2017), the impact of the corelease of DA and NE from LC axon terminal neurons on behavior identified here has large implications for the study of these two catecholamine neuromodulators across brain regions with dual VTA and LC innervation. Furthermore, understanding how these two LC-derived catecholamines function simultaneously to act on downstream GPCRs to influence neuronal activity will provide exciting insights into the fundamental neurobiology of neuromodulation and future pharmacotherapies for neuropsychiatric diseases.

## Abbreviations

LC: locus coeruleus
DA: dopamine
*Dbh*: dopamine beta hydroxylase
NE: norepinephrine
BLA: basolateral amygdala

## 4 METHODS

### Animals

Adult (20-35 g) male or female *Dbh*-cre or *Th*-cre mice and Cre (-) litter-mate control mice were used for *in vivo* experiments after backcrossing to C57BL/6J mice for at least 10 generations. Mice were group housed, given access to food pellets and water ad libitum, and maintained on a 12:12-hour dark/light cycle (lights off at 7:00 a.m.). Animals were held in a sound attenuated holding room facility in the lab starting at least one week prior to surgery, as well as post-surgery and throughout the duration of behavioral assays to minimize stress from transportation and disruption from foot traffic. All mice were handled and, where appropriate, connected to fiber optics two times a day for one week prior to behavioral experimental testing. All experimental procedures were approved by the Animal Care and Use Committee of Washington University and the Animal Care and Use Committee of University of Washington and conformed to NIH guidelines.

### Stereotaxic surgery

Surgeries were performed under 2% isoflurane anesthesia (Piramal Healthcare, Maharashtra, India). All injections in BLA and CA1 were done using a Hamilton blunt syringe (Reno, NV), while injections into the LC were completed using a Hamilton beveled syringe (Reno, NV).

For detecting NE and DA dynamics in ex-vivo slice experiments, Dbh-cre mice were injected with 500nL of AAV1-hSyn-GRAB_NE2m_-eGFP or AAV2/9-hSyn-WPRE-PA-GRAB_DA3m_-eGFP into dorsal CA1 (AP: 1.9, ML: −1.4, DV: −1.65) (Dbh-cre::BLA or CA1: GRAB_DA_ or GRAB_NE_). To activate terminals while recording, Chrimson was injected into the LC (AP: −5.45, ML: −1.10, DV: −4.3 to −3.9) of the same mice (Dbh-cre::LC:Chrimson). Mice were allowed to recover and viruses to express for a minimum of 5 weeks.

For recording initial neural dynamics in response to behaviors, either 500nL of AAV2/9-hSyn-WPRE-PA-GRAB_DA3m_-eGFP or AAV1-hSyn-GRAB_NE2m_ (both prepared in the UW NAPE center Viral Core, Seattle, WA) were injected with a Hamilton blunt syringe (Reno, NV) into the BLA (AP: −1.4, ML: −3.15, DV: −4.8 to −4.55), dCA1 (AP: 1.9, ML: −1.4, DV: −1.65) of Dbh-cre mice (Dbh-cre::BLA or CA1:GRAB_DA_ or GRAB_NE_). For investigating the neuromodulator release in response to optogenetic activation of terminals, Chrimson was injected with a Hamilton beveled syringe (Reno, NV) into the LC (AP: −5.45, ML: −1.10, DV: −4.3 to −3.9) of the same mice as those which we were detecting behavioral neural dynamics. After injections, fiber optic ferrules (Doric) were chronically implanted above either the BLA or dCA1, and dental cement (Lang Dental, Wheeling, IL) or C&B Metabond was used to secure the implants. BLA and dCA1 implants and injections were done in the same mice. Mice were allowed to recover and viruses to express for a minimum of 5 weeks.

For investigating the neuromodulator release in response to optogenetic activation of terminals, the red-shifted channelrhodopsin ChrimsonR was injected with a Hamilton beveled syringe (Reno, NV) into the LC (AP: −5.45, ML: −1.10, DV: −4.3 to −3.9) of Th-cre mice (Th-cre::LC:Chrimson). To inhibit the VTA, the chemogenetic inhibitor AAV5-hSyn-DIO-HA-hM4D(Gi) was injected bilaterally in the VTA of the same Th-cre mice (Th-cre::VTA^bilat^:Gi-DREADDs). For detecting NE and DA dynamics in the same Th-cre mice, 500nL of AAV1-hSyn-GRAB_NE2m_-eGFP or AAV2/9-hSyn-WPRE-PA-GRAB_DA3m_-eGFP were injected into dorsal CA1 (AP: 1.9, ML: −1.4, DV: −1.65) and the BLA (AP: −1.4, ML: −3.15, DV: −4.8 to −4.55) (Th-cre::BLA or CA1:GRAB_DA_ or GRAB_NE_). After injections, fiber optic ferrules (Doric) were chronically implanted above the dCA1 (AP: 1.9, ML: −1.4, DV: −1.65) and the BLA (AP: −1.4, ML: −3.15, DV: −4.8 to −4.55), and C&B Metabond was used to secure the implants. We waited five weeks for viruses to express before initiation of stimulation experiments.

For dual-color neuromodulator recordings, 500nL of AAV9-hSyn-GRAB_rDA2m_ and AAV1-hSyn-GRAB_NE2m_-eGFP were injected into dorsal CA1 (AP: 1.9, ML: −1.4, DV: −1.65) and the BLA (AP: −1.4, ML: −3.15, DV: −4.8 to −4.55) (Th-cre::BLA or CA1:GRAB_rDA_ and GRAB_NE_). We injected the cre-dependent chemogenetic inhibitor AAV5-hSyn1-DIO-HA-hM4D(Gi) bilaterally in the LC or VTA of Th-cre mice (Th-cre::LC^bilat^:Gi-DREADDs) or (Th-cre::VTA^bilat^:Gi-DREADDs). After injections, fiber optic ferrules (Doric) were chronically implanted above the dCA1 (AP: 1.9, ML: −1.4, DV: −1.65), and C&B Metabond was used to secure the implants. We waited five weeks for viruses to express before initiation of stimulation experiments.

### Tissue collection and immunohistochemistry

After the conclusion of behavioral testing, mice were anesthetized with sodium pentobarbital and transcardially perfused with ice-cold PBS, followed by 4% phosphate-buffered paraformaldehyde. Brains were removed, postfixed overnight in 4% paraformaldehyde, and then saturated in 30% phosphate-buffered sucrose for 2–4 days at 4°C. Brains were sectioned at 30mM on a microtome and stored in a 0.01M phosphate buffer at 4°C prior to immunohistochemistry and tracing experiments. For behavioral cohorts, viral expression and optical fiber placements were confirmed before inclusion in the presented datasets. Viral expression and implant placements were verified using fluorescence and confocal (Leica Microsystems) microscopy. Images were produced with 10X, 20X, and 63X objectives and analyzed using ImageJ software (NIH) and Leica Application Suite Advanced Fluorescence software.

### Looming stimulus

Adult mice were put in a custom-made open clear acrylic box, (50cm x 50cm) within a sound-attenuated room maintained at 23°C. The open box had white paper on all walls to limit outside stimuli. Mice were allowed to roam the box for 11 minutes prior to stimulus presentation to habituate to the box. At 11 minutes and in 60s intervals, a looming posterboard blocking overhead lighting was held over the arena for 10s. This was repeated 5 times. Fiber photometry was recorded for multiple biosensors detecting neural dynamics.

### Reward retrieval and consumption

Mice were initially food deprived to 90% of their body weight. Prior to testing, we put multiple sucrose pellets per mouse in their home cage to extinguish neophobia of this reward. On testing days, we placed mice in a modular (17.8×15.2×18.4cm^3^; Med Associates Inc.) arena where they were allowed to freely roam. Using MEDPC program, we randomly delivered rewards at a pseudorandom ITI of 90-150s, with a total of 18 possible rewards in each session. Fiber photometry was recorded for multiple biosensors detecting neural dynamics.

### Shock stimulus

Mice were placed in a modular arena and allowed to freely roam. After a baseline period of 180s in the arena, mice experienced the shock of a 0.5-0.7mA current run through the metal grating floor of the arena for 2s. There was a pseudorandom ITI of 20-90s, with 10 total shocks. Fiber photometry was recorded for multiple biosensors detecting neural dynamics. Video was recorded for the duration of the shock protocol and for 180s after. EzTrack(Pennington et al., 2019) was used to track and align freezing bouts for behavioral analysis.

### *In vivo* fiber photometry

An optic fiber was secured to implanted fibers on experimental mice with a ferrule sleeve (Doric, ZR 2.5). Two light-emitting diodes (LEDs) were driven by a real-time processor (TDT, either RZ5P or RZ10x) that recorded dynamics from optic fibers detecting biosensor activity expressed in the brain, with 531-Hz sinusoidal 470nm (465nm on the RZ5P) and 211-Hz sinusoidal 405nm light. The 470nm LED was used to record emitted light from green sensors (GRAB_NE_, dLight, and GRAB_DA_) while the 405nm LED served as an isosbestic control. In dual sensor recordings taken with the RZ10x, 311-Hz sinusoidal 535nm light was used to record red sensors (GRAB_rDA_). LED power for each LED wavelength were measured at the tip and adjusted to 30uW before each day of recording. For optogenetic stimulation, a 625nm LED was pulsed with 5ms pulse width for 3 or 30s. Laser power was adjusted to obtain 15mW transmittance into the brain.

### Photometry analysis

A custom MATLAB script was developed for analysis of photometry recordings in the context of mouse behavior. For each photometry recording, a linear-least squares regression was used to find the projection of the isosbestic curve onto the 470 nm excitation photometry trace. The best fit isosbestic 405 nm excitation control signal was subtracted from the 470nm excitation signal to remove artifacts from intracellular NE-dependent sensor fluorescence(Lerner et al., 2015). Baseline drift was evident in the signal due to slow photobleaching artifacts, particularly the first several minutes of each recording session, and the linear least squares subtraction of the isosbestic control signal eliminated the drift seen in the 470nm excitation signal. After centering the signal around the first three minutes of the session, the photometry trace was z-scored relative to the mean and standard deviation of the test session. The post-processed fiber photometry signal was analyzed by time-locking each onset of behavior or stimulus, centering the data, and averaging the signal.

### *Ex vivo* slice preparation

Mice were briefly anesthetized with isofluorane. Mice were perfused with cold N-methyl-d-glucamine (NMDG)–substituted artificial cerebrospinal fluid (aCSF) containing 93mM NMDG, 2.5mM KCl, 1.25mM NaH2PO4, 30mM NaHCO3, 20mM Hepes, 25mM glucose, 5mM ascorbic acid, 2mM thiourea, 3mM Na-pyruvate, 12mM N-acetyl-l-cysteine, 10mM MgSO4, and 0.5mM CaCl2. After perfusion, brains were extracted. Coronal sections of the hippocampus were cut to a slice thickness of 250μm using a vibrating Leica VT1000S microtome in a bath of cold, oxygenated NMDG solution. The slices were then placed in hot NMDG aCSF solution (32-34 degrees C) for 10min. Then slices were placed in a room temperature holding chamber filled with HEPES solution consisting of: 92mM NaCl, 2.5mM KCl, 1.2mM NaH2PO4, 30mM NaHCO3, 20mM Hepes, 25mM glucose, 2mM CaCl2, 2mM MgCl2 (pH adjusted to 7.3 to 7.4 with NaOH). The samples were kept in the HEPES solution for at least 30min prior to 2-photon imaging and were covered to minimize photobleaching. For both HEPES and NMDG osmolality was confirmed to be between 300-310mOsm/kg, and pH was confirmed to be between 7.3-7.4.

### 2-photon slice imaging, optogenetic stimulation, and pharmacology

Flow rate of ACSF or drug was maintained at 2mL/min. Temperature of flowing ACSF was controlled at 31-34 degrees C by the temperature controller TC-324C before any slice was added to the bath. An anchor was used to keep the brain slice stable while ACSF or ACSF and drug was washed on. A vacuum pump ensured a steady 2mL/s of flow. An Olympus 4x UPlanFL N lens was used to localize expression, and then an Olympus 20x XLPLN25XWMP lens 2mm working distance for water immersion, and we acquired images with Olympus FV31S-SW software. The 2-photon scope used was an FVMPE-RS Olympus 2p scope powered by a Mai Tai Tisapphire laser with dispersion compensation running at 920nm. Individual frames were acquired at 0.92hz using a Galvano scanner with a resolution of 512px x 512px. After setting up a slice on the 2-photon rig, we ran a positive control trial by washing high concentration (100μM) reactive ligand over the sensor-expressing slice. Once reactivity to the reactive ligand was confirmed, we allowed oxygenated ACSF to flow for 5 minutes for ligand to unbind from sensor. We next initiated optogenetic stimulation of brain slices using a 615nm red LED (CoolLED) controlled by a master-9 stimulator, programmed for either 5hz or 20hz stimulation for 3s or 30s, with a pulse-width of 5ms and limited to power of 3mW. Ligand solutions were oxygenated for at least 5min prior to washing on slices. Ligands were prepared with concentrations of between 3nM to 100μM.

Following optogenetic stimulation trials, we allowed oxygenated ACSF to flow over slices for at least 5 minutes for ligand to unbind from sensors before initiating ligand washes for concentration response curves. In some washes, we pumped ligand for 60s and recorded for two additional minutes to capture sensor binding and unbinding. We waited one minute between sessions of low concentration and 3-5min for higher concentration washes. We confirmed fluorescence resembled baseline of the previous session before initiating another wash, and would alternate between high concentration ligand washes and low concentration washes (30μM, 10nM, 10μM, etc).

### Pupillometry

Mice were anesthetized with 2% isofluorane. An infrared light was set up to record the mouse’s pupil from a FLIR camera (1:1.4CCTV Lens, 9-22 mm lens with 1/3” focal length). Pupil contrast was carefully checked before recording session started. After 10min of recording for a baseline, we injected mice with either CNO (0.5 mg/kg i.p.) or saline vehicle. At least 45min after injection were recorded to track any changes in radii due to inhibition of the LC. We used a pupillometry analysis toolbox developed and described in Privitera et al., 2020.

### DBH-KO mice experiments

A total of 40 Dbh(−/−)mice, maintained on a mixed 129/SvEv and C57BL/6 J background as previously described(Thomas et al., 1995; Thomas and Palmiter, 1997), were used in this study. *Dbh*(−/−) males were bred to *Dbh*(+/−) females, and pregnant *Dbh*(+/−) dams were administered drinking water containing the β-adrenergic receptor (AR) agonist isoproterenol and the α1AR agonist phenylephrine (20 μg/ml each; Sigma-Aldrich) with vitamin C (2 mg/ml) from E9.5E14.5, and the synthetic NE precursor L-3,4-dihydroxyphenylserine (DOPS; 2 mg/ml; Lundbeck, Deerfield, IL) + vitamin C (2 mg/ml) from E14.5-parturition to prevent embryonic lethality resulting from complete *Dbh* deficiency. *Dbh*(−/−) mice are easily distinguished from their NE-competent littermates by their visible delayed growth and bilateral ptosis phenotypes. *Dbh*(+/™) littermates were used as controls because their behavior and catecholamine levels 5 are indistinguishable from wild-type *Dbh*(+/+) mice(Bourdélat-Parks et al., 2005; Szot et al., 1999; Thomas and Palmiter, 1997). *Dbh*(−/−) males were bred to *Th*-cre females to produce *Th*-cre *Dbh*(−/−) mice. All animal procedures and protocols were approved by the Emory University Animal Care and Use Committee, in accordance with the National Institutes of Health guidelines for the care and use of laboratory animals. Mice were maintained on a 12 h/12 h light/dark cycle (7:00/19:00 h) with access to food and water ad libitum, except during behavioral testing. Behavioral testing was conducted under standard lighting conditions during the light cycle (14:00-17:00) in the same room where the mice were housed to minimize the stress of cage transport on test days. To record NE or DA dynamics, 500nL of AAV1-hSyn-GRAB_NE2m_ (prepared in the UW NAPE center Viral Core, Seattle, WA) or AAV2/9-hSyn-GRAB_DA3m_-WPRE-PA (Brain VTA) were injected with a Hamilton blunt syringe (Reno, NV) into CA1 (AP: 1.9, ML: −1.4, DV: −1.65) of *Dbh*(+/+)-cre, *Th*-cre *Dbh*(+/−), and *Th*-cre *Dbh*(−/−) mice. For inhibition experiments, AAV5-hSyn1-DIO-HA-hM4D (Gi) were injected bilaterally in the LC (AP: −5.45, ML: −1.10, DV: −4.3 to −3.9) or the VTA (15 degree angle, AP: −3.25, ML: −1.6, DV: −4.7 to −4.2). We waited five weeks for viruses to express before initiation of aversive and appetitive experiments. For investigating the neuromodulator release in response to optogenetic activation of terminals, the red-shifted channelrhodopsin ChrimsonR was injected with a Hamilton beveled syringe (Reno, NV) into the LC (AP: −5.45, ML: −1.10, DV: −4.3 to −3.9). After injections, fiber optic ferrules (Doric) were chronically implanted above the dCA1 (AP: 1.9, ML: −1.4, DV: −1.65), and C&B Metabond was used to secure the implants. We waited five weeks for viruses to express before initiation of stimulation experiments, done in accordance with photometry methods.

### Quantification and statistical analysis

All data are expressed as mean ± SEM. Behavioral data were analyzed with GraphPad Prism 10.0 (GraphPad, La Jolla, CA). Two-tailed student’s t test, one-way or two-way ANOVAs were used to analyze between-subjects designs. Repeated-measures designs were analyzed using a mixed-effects restricted maximum likelihood (REML) model. Tukey was used for post-hoc pairwise comparisons. The null hypothesis was rejected at the p < 0.05 level. Statistical significance was taken as ^*^ p < 0.05, ^**^ p < 0.01, ^***^ p < 0.005, n.s. represents not significant. All statistical information is listed in Supplementary Table 1.

## Acknowledgements

We thank Dr. Azra Suko for lab management and organization and Carina Pizzano, Akshay Rana, Bailey Wells, and Valerie Lau for assistance with the mouse colony. We thank Dr. Paul Phillips for feedback on sensor and genetic knockout experiments. We also thank the entire Bruchas laboratory as well as the Neuroscience of Addiction, Pain, and Emotion (NAPE) Center at the University of Washington for resources alongside critical feedback.

## SUPPORTING INFORMATION

Additional supporting information may be found in the online version of the article.

## SUPPLEMENTAL FIGURES

**Supplemental Data Figure 1:**
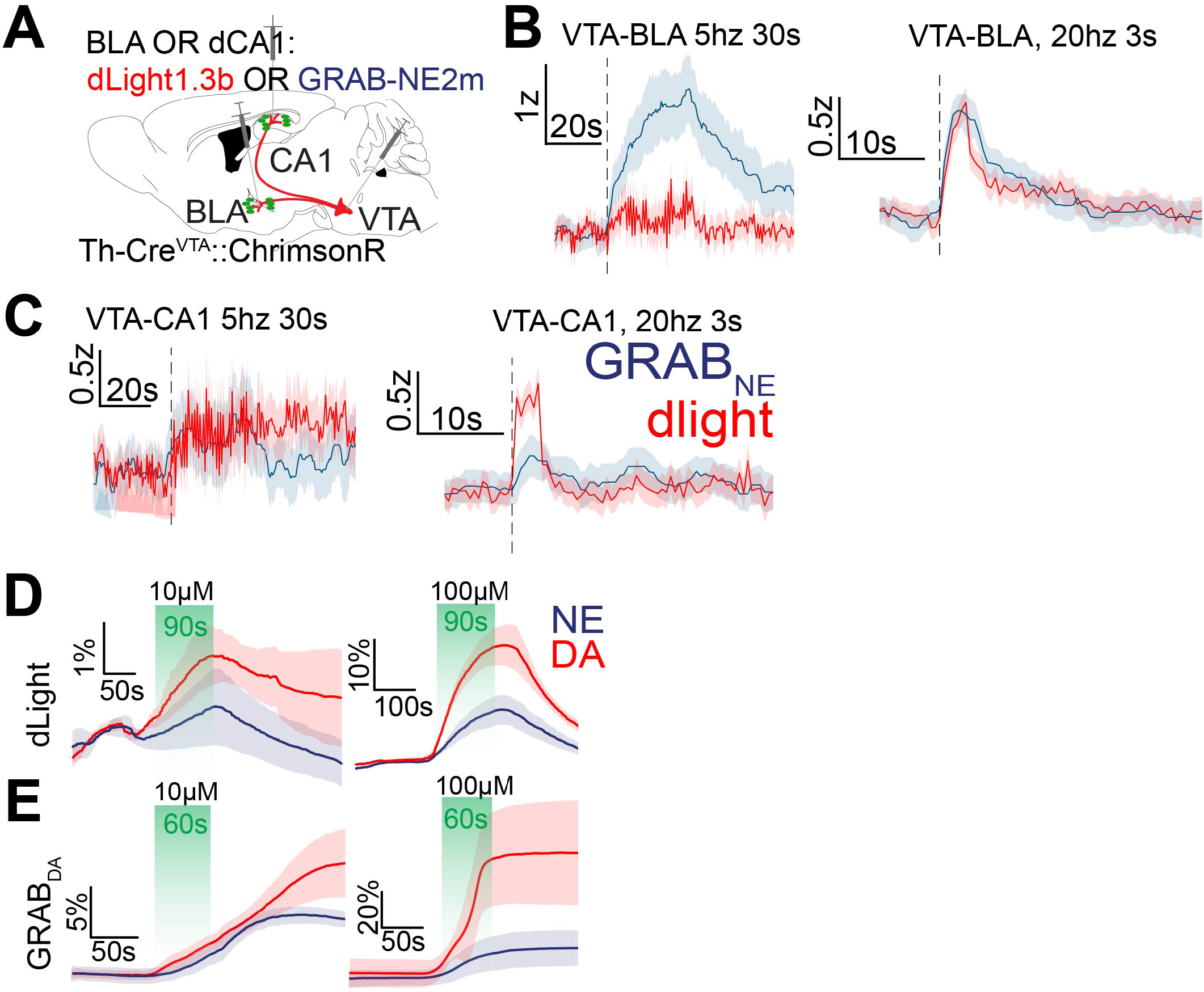
Controls for biosensor selectivity. (a) Schematic of experimental approach of surgical injections for VTA terminal stimulation and recording photometric signals. Schematic depicts infection of VTA-DA cells with ChrimsonR (AAV5-hSyn-FLEX-ChrimsonR-tdTomato) and GRABNE or dLight in BLA (b) or CA1 (c). (b-c) Stimulation of VTA terminals in BLA (b) and CA1 (c), at 5hz for 30s (left) or 20hz for 3s (right). D-E. ΔF/F% change during DA and NE washes on dlight (G) or GRABDA (H) expressing slices at either 10μM (left) or 100μM (right).

**Supplemental Data Figure 2:**
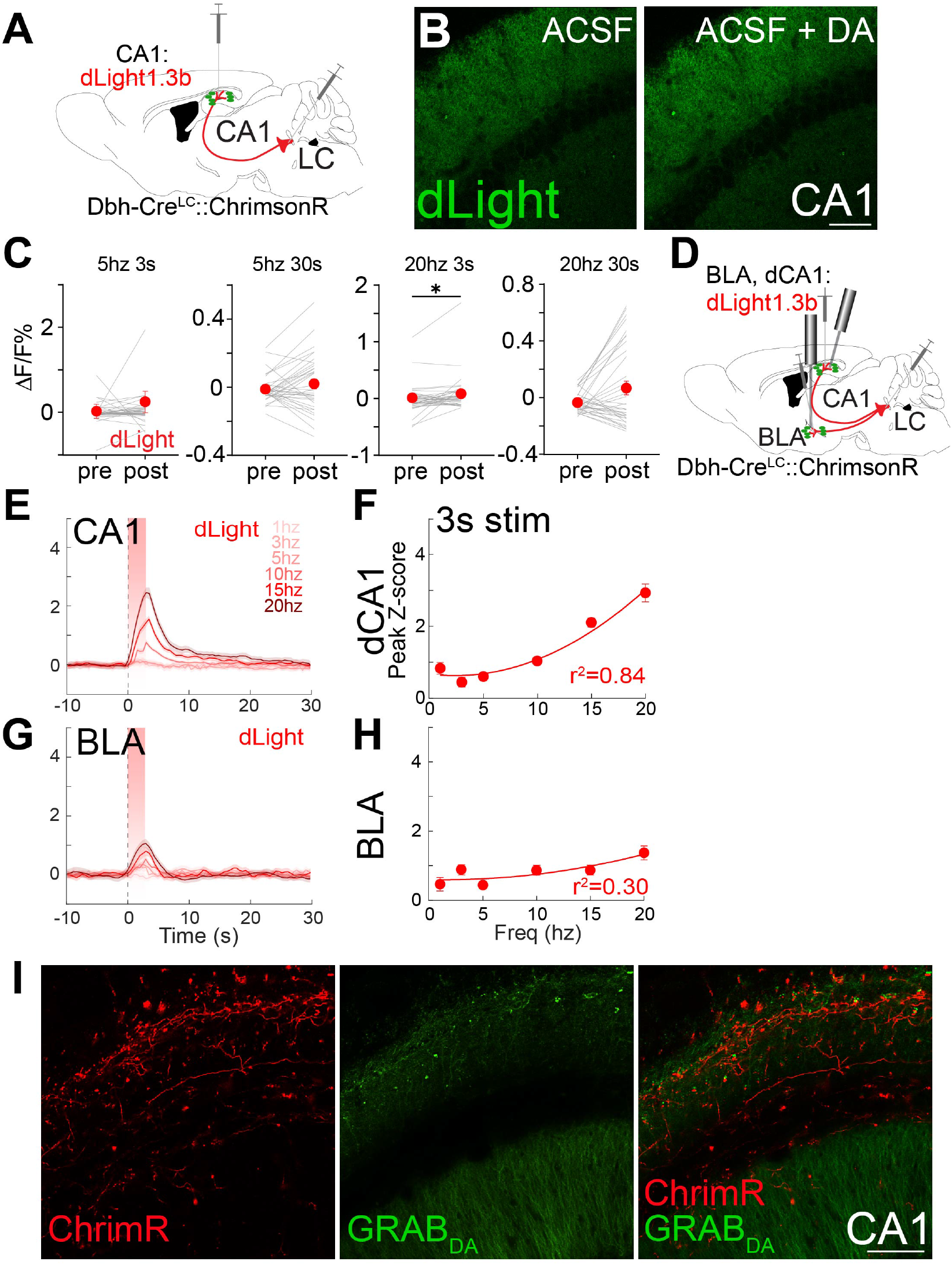
LC-evoked DA is detected by dLight *in vivo* and *ex vivo*. (a) Schematic of surgeries for 2-photon experiments with dLight. Chrimson was expressed in LC, and dLight in CA1. (b) Vehicle (left) and 30μM DA (right) washes on dLight expressing slices. Scale bar, 100μm. (c) ΔF/F% change from before to after stim in dlight (n=4 biological replicates, 6 slices) in response to a range of stimulation paradigms: 5hz 3s, 5hz 30s, 20hz 3s, 20hz 30s (left to right). Wilcoxon test, p=0.027. (d) Surgery schematic for in vivo optogenetic stimulation, with dLight expressed in CA1 and BLA and Chrimson expressed in LC. (e) dLight response to LC-CA1 stimulation at a range of frequencies (1-20hz). (f) Frequency response curves of dLight response when stimulating LC-CA1 terminals for 3s. Second-order curve fit to dLight, n=5-7, r^2^= 0.84. (g) dLight response to LC-BLA stimulation at a range of frequencies (1-20hz). (h) Frequency response curves of dLight response when stimulating LC-BLA terminals for 3s. (i) Representative 2-photon image of virus expression in a hippocampal slice expressing GRABDA in CA1 and ChrimsonR in LC terminals in CA1. Scale bar, 100μm.

**Supplemental Data Figure 3:**
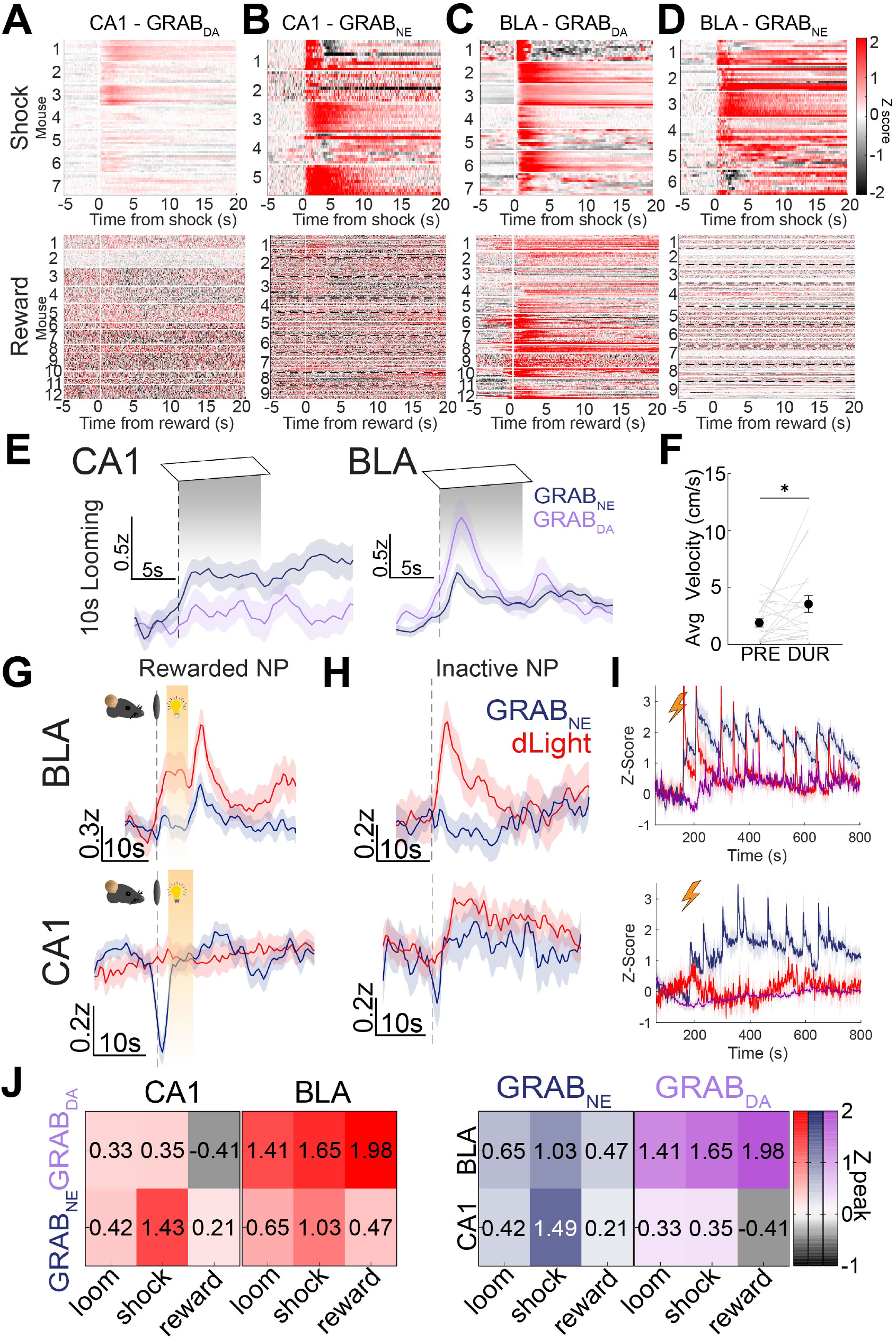
NE and DA dynamics during aversive and appetitive behaviors. (a-d) Heatmaps of GRABDA and GRABNE response to shock (top) and reward (bottom) in BLA (a: GRABDA in CA1, b: GRABNE in CA1). (c-d) Heatmaps of GRABDA and GRABNE response to shock (top) and reward (bottom) in BLA (c: GRABDA in BLA, d: GRABNE in BLA). (e) NE (blue) and DA (purple) response to looming in CA1 (left) and BLA (right). (f) Mice dart at the first three looming stimuli presentations. Paired t-test, p=0.0364. (g-h) Response to nosepoking in the rewarded noseport (g), the unrewarded noseport (h) for BLA (top) or CA1 (bottom). (i) Full session recording of GRABDA (purple), GRABNE (blue), or dLight (red) during shock in BLA (top) or CA1 (bottom). (j) Heatmap of peak z-score in response to looming stimuli, shock, reward in CA1 and BLA.

**Supplemental Data Figure 4:**
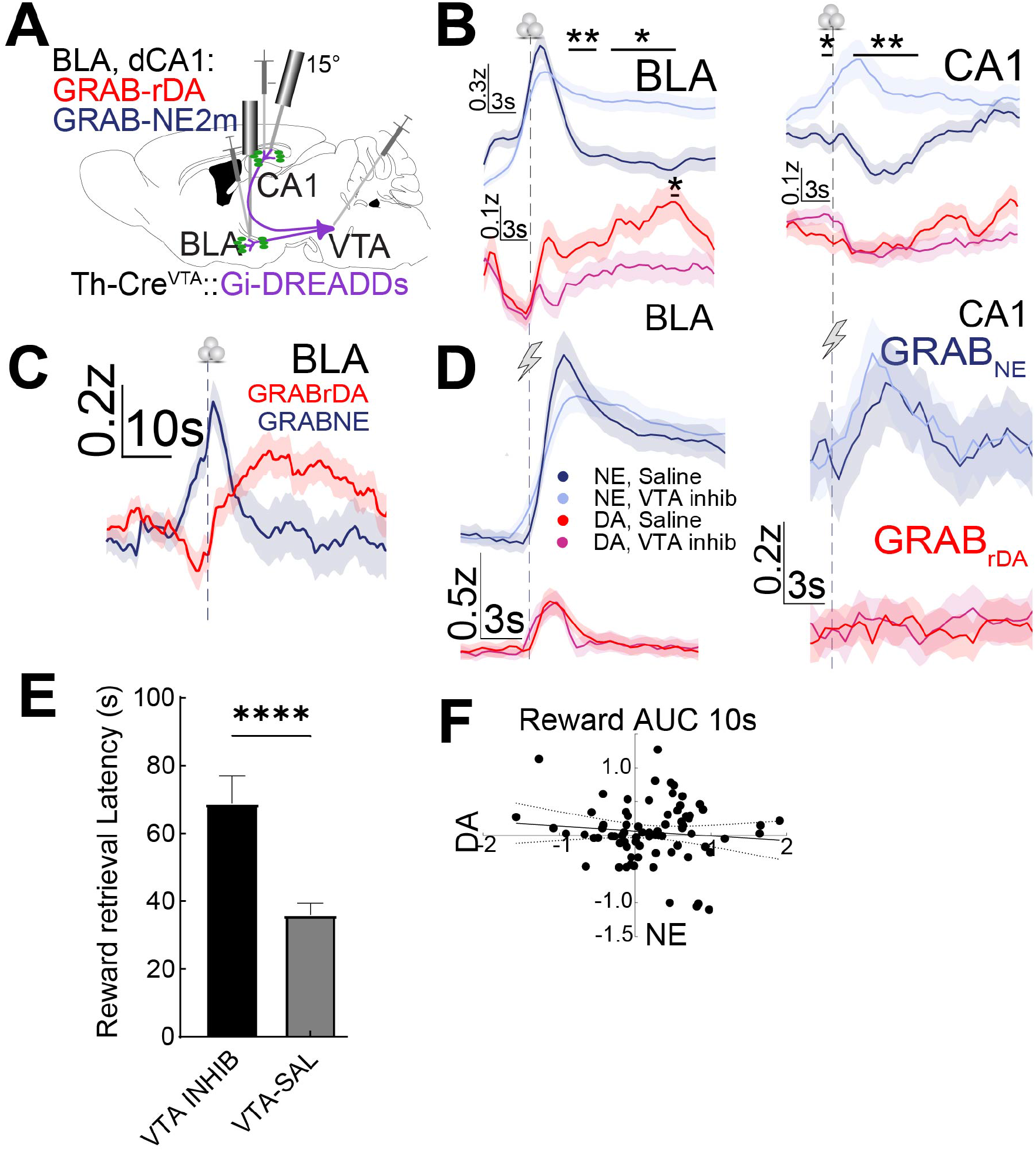
VTA or LC inhibition effects on DA release in reward and shock. (a) Schematic of surgeries preparing for recording DA and NE dynamics while inhibiting VTA. (b) DA response to reward in BLA (left) or CA1 (right) under VTA inhibition (magenta) or vehicle (red). (c) Latency to retrieve rewards following either saline or CNO administration, in mice expressing hM4Di in VTA. (d) Averaged, simultaneously recorded GRAB_rDA_ and GRAB_NE_ dynamics in BLA during reward retrieval prior to any neuromodulation. (e) AUC for 10s of DA vs AUC for 10s of NE after reward retrieval in Fig S4D. R^2^=0.01. (f) DA response to shock in BLA (left) or CA1 (right) under VTA inhibition (magenta) or vehicle (red). Significance of 1s bins for all traces calculated using 2-way ANOVA with Sidak test for multiple comparisons, p<0.05.

